# Variation in CURE instruction and limitations of CURE studies undermine what can be concluded about CURE effects on student outcomes

**DOI:** 10.64898/2026.08.04.741794

**Authors:** W. Nathan Lantz, Erin L. Dolan, Swara Barvalia, Sydney Apraku, Harika Kandru, Madhumitha Senthamizh Selvan, Ania A. Majewska

## Abstract

Research suggests that students realize a range of cognitive, affective, and motivation-related outcomes by participating in course-based undergraduate research experiences (CUREs). However, the overall effects of CUREs on student outcomes are unclear. We sought to address this knowledge gap by embarking on a meta-analysis of CURE studies using the Population, Intervention, Comparison/Control group, and Outcomes framework. We conducted a literature search to produce an analytic sample of 353 studies of the effects of CUREs instruction on undergraduates. We characterized the CUREs being studied in this sample, including their dose, duration, and design. We also examined each study to determine if outcomes were measured before and after the CURE and with a comparison or control group so that effects could be attributed to CURE. We found substantial limitations in studies of CUREs, which precluded meta-analysis. CUREs varied widely and were unevenly described, making it difficult to determine what could or should constitute CURE instruction as an intervention. Many outcomes were studied, often without a comparison or control group and with inconsistent measurement and reporting. Our results indicate that advancements are needed in the design and reporting of research on CUREs to gauge their effects on student outcomes.

**Highlight Summary:** Many studies of CURE instruction aim to show effects on student outcomes. Yet, systematic review of these studies reveals limitations in study design, methods, and reporting that preclude meta-analysis and limit any cross-study conclusions.

## INTRODUCTION

Course-based undergraduate research experiences (CURE) engage students enrolled in a course in conducting research that is of interest beyond the classroom rather than completing lab exercises with predictable results (Auchincloss et al., 2014; Cooper et al., 2019). The potential of CUREs to increase access to research opportunities, especially in STEM, has led to the growth of this instructional approach across the U.S. (National Academies of Sciences, Engineering, and Medicine, 2015). This is evident in the growing number of CURE publications over the past decade (Beck et al., 2023; Buchanan & Fisher, 2022; Treibergs et al., 2025). As CUREs have become more widespread, so have efforts to understand the effects of CURE instruction on student experiences and outcomes. Recent reviews of CURE literature have revealed that CUREs are being developed, taught, and assessed at diverse institutions, in a range of disciplines, and for a variety of student populations (Buchanan & Fisher, 2022; Krim et al., 2019; Watts & Rodriguez, 2023). This growth in research on CUREs presents an opportunity to understand the effects of CUREs on student outcomes.

Research suggests that CURE students can realize a range of cognitive, affective, and motivation-related outcomes. Some studies indicate that participating in a CURE can enhance students’ knowledge and skills, including their knowledge of scientific concepts (e.g., Shaffer et al., (2010), understanding of the nature of science (e.g., Russell & Weaver, 2008), and their skills in thinking critically and from a systems perspective (e.g., Carson, 2015; Stanfield et al., 2022). Other studies explore the influence of CURE instruction on students’ attitudes, beliefs, and dispositions, such as their scientific self-efficacy (e.g., Olimpo et al., 2016), beliefs about scientific research process (e.g., Goodwin et al., 2021) and interest in science (e.g., DeChenne-Peters et al., 2023). Participation in CUREs has been associated with shifts in students’ identification as scientists (e.g., Esparza et al., 2020) and their sense of belonging in the scientific community (e.g., Merkle et al., 2023). Participation in CUREs has also been causally linked to higher rates of retention in STEM majors and increased graduation rates (Hanauer, Graham, SEA-PHAGES, et al., 2017; Rodenbusch et al., 2016). Collectively, these studies have spanned institution types (e.g., community colleges, comprehensive universities, research intensive universities), suggesting that the benefits of CUREs may be applicable to students across institutional contexts. When considered together, these results suggest that there is a sufficient body of research on CURE instruction to determine the effects of CUREs for students.

Understanding student outcomes stemming from CUREs requires a systematic approach to synthesizing published work. Meta-analysis offers a powerful method for this synthesis (Maynard, 2024; Pellegrini et al., 2025). A meta-analysis is a statistical method that combines results from multiple independent but similar studies to calculate an overall effect. Meta-analysis enables researchers to determine the size of an overall effect represented in an accumulated body of evidence. Meta-analysis can also be useful for characterizing the degree and sources of heterogeneity in the data across multiple studies (Pigott & Polanin, 2020). This method is particularly valuable in education, where individual studies vary in sample sizes, design, and context, making it difficult to draw conclusions from a single study that cannot fully represent the student population. Thus, meta-analysis offers a way to determine the average effect of an instructional approach, such as CURE, across studies, and enhance understanding of the factors that influence the size of the effect. Furthermore, findings from meta-analyses can inform practical recommendations for implementation of an intervention, such as CUREs. Here we sought to conduct a meta-analysis of CURE studies to understand CURE effects on student outcomes.

### Meta-analytic framework

Given our goal to conduct a meta-analysis of CURE studies, we designed our study using the Population, Intervention, Comparison, Outcome (PICO) framework, a meta-analytic approach well-suited for charactering the effects of an intervention. PICO was designed to promote evidence-based medicine by enabling systematic reviews and meta-analyses of quantitative research to inform clinical practice (Chandler et al., 2019; Richardson et al., 1995). PICO supports researchers in conducting systematic reviews and meta-analysis that clearly identify the group of interest (*<u>P</u>opulation*), the treatment, exposure, or procedure evaluated (*Intervention*), what the intervention is being measured against (*<u>C</u>omparison* or *<u>C</u>ontrol* groups), and what is being measured (*<u>O</u>utcome*). As such, PICO guides literature search and inclusion criteria (Chandler et al., 2019). For instance, for a meta-analysis of CUREs, the *Population* would be undergraduate students, the *Intervention* would be the educational experience (or range of experiences) that counts as a CURE, the *Comparison or Control* would be the undergraduate students and/or educational experiences used as a reasonable standard for comparing outcomes, and the *Outcome* would be the reported effect(s) of CURE instruction on undergraduate student outcomes, including sample sizes, means, standard deviations, and effect sizes (e.g., t-statistic), or data to calculate them (Pigott & Polanin, 2020).

Other meta-analytic frameworks have been proposed to better accommodate qualitative research, including qualitative findings from mixed methods studies. For example, the SPIDER framework (Sample, Phenomenon of Interest, Design, Evaluation, Research type) was developed to enable systematic reviews that are inclusive of a variety of study designs (Cooke et al., 2012). Side-by-side testing of PICO and SPIDER in search methods revealed that the PICO approach was more sensitive to finding relevant literature (Methley et al., 2014). In addition, PICO is better suited to enabling cross-study quantitative analysis while SPIDER is better suited to cross-study synthesis of results, including qualitative findings. Given the variety of how CUREs might be described and studied in the literature and our interest in determining the strength of any effects of CURE instruction, we opted to use PICO to structure our meta-analysis and to focus on quantitative findings.

### Current study

We sought to conduct a meta-analysis of CURE literature to determine the overall strength of the relationship between CURE instruction and student outcomes. We used the PICO framework to attempt to identify a suitable body of literature. Although identifying the population as undergraduate students was straightforward, we also needed to define the intervention, comparison or control groups, and outcomes. To accomplish this, we addressed the following questions:

1. To what extent do CURE studies describe the instructional experience sufficiently to clearly and consistently identify the **intervention**?
2. To what extent do CURE studies include **comparison or control groups**?
3. What student **outcomes** are being studied for CUREs, and to what extent are these outcomes being measured and reported in ways that enable meta-analysis?

Ultimately, we determined that the CURE literature was insufficient to conduct a meta-analysis. However, by addressing these questions, we discovered several noteworthy trends in research on CUREs, which we report here. We also offer recommendations for future research on CURE instruction based on our results.

## METHODS

Here we describe our methods for conducting our CURE literature search, review, and analysis in terms of the PICO framework (*Population*, *Intervention*, *Comparison or Control*, and *Outcomes*). We followed the Preferred Reporting Items for Systematic Reviews and Meta-Analyses (PRISMA) checklist (Page et al., 2021), which provides guidelines for conducting and reporting the methods used to review literature (see Figure 1 for a visual overview). We used the web platform Covidence (https://covidence.org) to manage the process of reviewing literature and applying our inclusion and exclusion criteria. We used R programming version 4.5.3 for calculating descriptive statistics and visualizing results (R Core Team, 2026).

**Figure 1.**
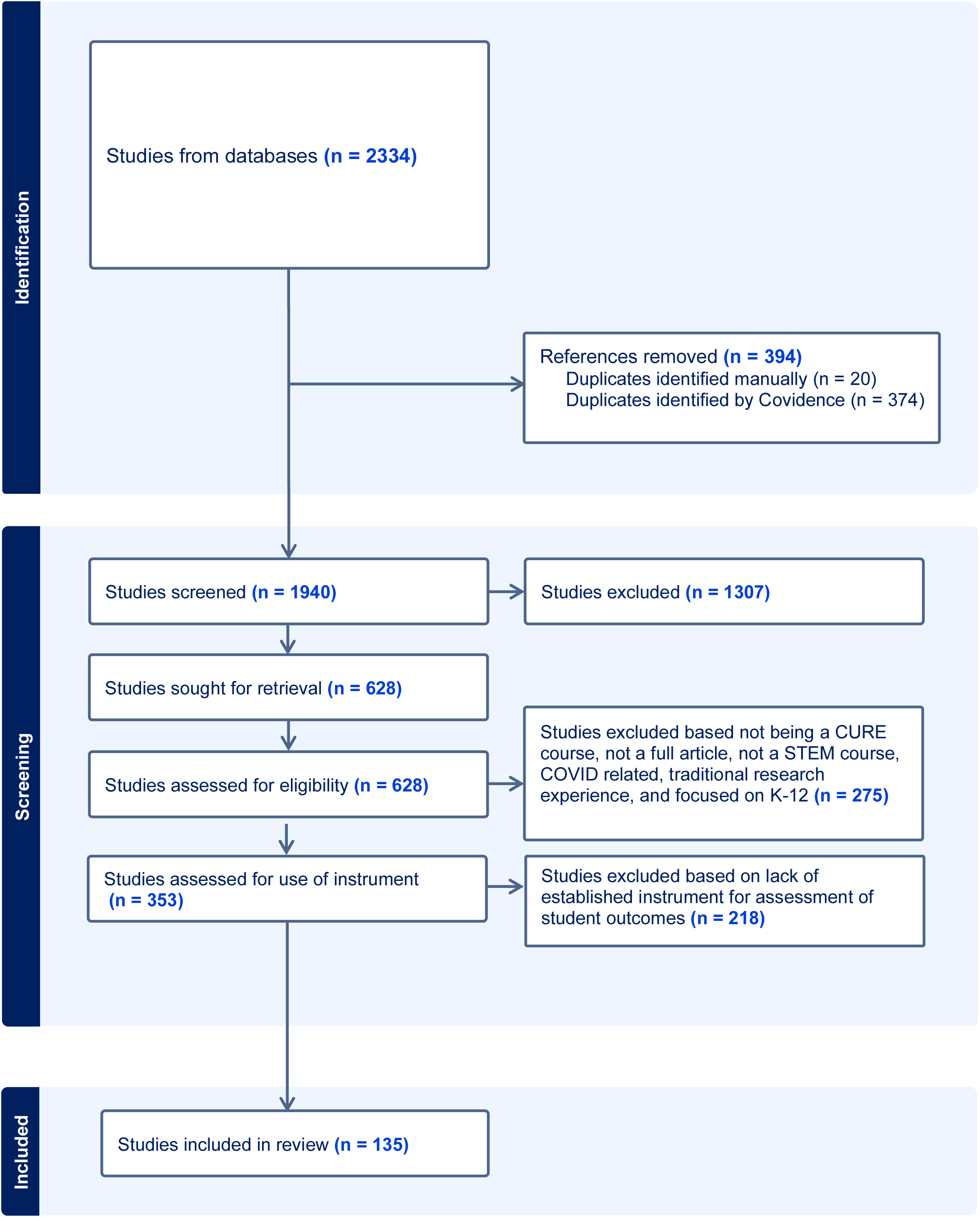
PRISMA flow diagram specifying the number of studies identified for initial review, removal of duplicates, and screening for inclusion in our dataset.

### Systematic literature search

We started by selecting search terms intended to identify literature related to our *Population* (i.e., undergraduate students) and *Intervention* of interest (i.e., CURE instruction). Specifically, we used the following terms to capture literature describing credit-bearing courses in which undergraduate students conducted research with unknown outcomes that are of interest beyond the course: “course-based undergraduate research” OR “authentic research experience” OR “CURE” OR “class-based research” OR “research-based course” OR “discovery-based course.” We searched for published studies in Google Scholar, PubMed, ERIC, and Web of Science. The search resulted in 2,334 papers published between January 2002 and February 2023, which we imported to Covidence. Covidence removed 374 duplicates, resulting in 1,960 papers. We identified 20 additional duplicates during initial screening (described next), reducing our paper sample to 1,940.

### Inclusion and exclusion process

Two reviewers performed initial title and abstract screening in Covidence to identify whether each of the 1,940 papers met inclusion and exclusion criteria. During this screening, we retained literature that met five inclusion criteria: (1) peer-reviewed papers, (2) written in English, (3) full-text available online, (4) describing undergraduate students participating in a STEM CURE, and (5) examining and reporting student outcomes following the CURE. We excluded books, preprints, datasets, theses and dissertations, and conference abstracts, as well as studies that did not explicitly examine student outcomes following a CURE or that investigated non-CURE courses or non-STEM courses. We excluded studies that examined CURE outcomes during the COVID-19 pandemic given the potential for confounding effects on instruction. We also excluded studies of students in secondary education as outside of our population of interest and summer research experiences as outside of our intervention of interest. The title and abstract screening resulted in 628 papers.

We then proceeded with a full-text review of these 628 papers to determine they fully met our inclusion criteria. Each paper was reviewed by at least two members of the research team. If they did not agree on inclusion/exclusion, a third researcher reviewed the paper and made a final decision. During full-text review, we excluded 275 papers for not referencing a CURE-type experience in STEM, for not being a full article or primary literature (e.g., abstract, book, preprint, theses), for describing a course moved online due to the COVID-19 pandemic, or for focusing on K-12 student populations. Full-text review produced 353 articles that met our initial inclusion and exclusion criteria.

Finally, we sought to identify the subset of the 353 papers describing measurements of student *Outcomes* using a published instrument (e.g., survey, test). Our rationale for focusing on papers with published instruments was that we could review how the instrument was developed and what it was intended to measure and thus have some confidence about the validity of any inferences made using the instrument. To accomplish this, we employed Elicit (elicit.com), an artificial intelligence (AI) tool to assist with determining which studies cited published instruments that could have been used to measure student outcomes. Elicit works by converting PDFs into text and employing a large language model and natural language processing to analyze the text and report information based on user instructions (see Table S1 for our instructions for Elicit). We prompted Elicit to extract the “assessment instruments” from the 353 studies, which produced a list of 48 instruments. We then prompted Elicit to query studies and report whether each instrument was present in the study. We reviewed the Elicit results alongside the full text of each study to confirm that a given instrument was used to assess student outcomes in the publication (versus just being cited but not used or used but with no data reported). The process resulted in 135 studies that met our final inclusion criterion: reporting outcomes measured using a published instrument. This set of papers comprised our final analytic sample.

### Intervention description

To address our first research question (*To what extent do CURE studies describe the instructional experience sufficiently to clearly and consistently identify the intervention?*), we leveraged Elicit to conduct a preliminary characterization of the CUREs being studied in our analytic sample, including their dose, duration, and design. We prompted Elicit to review each paper and extract information about duration of the CURE described in the paper (i.e., number of semesters or weeks) and its dose (i.e., contact hours per week). If Elicit could not identify this information, it reported that the information was “not specified.”

Then, we used Elicit to provide a preliminary assessment of whether each study described the design of the CURE (see Table S1 for Elicit variable instructions). Specifically, Auchincloss and colleagues (2014) proposed that CUREs have five features that make them distinctive from other lab learning experiences – that students: 1) use of *scientific practices*, 2) have the potential to make *discoveries*, 3) do work that is *relevant* to stakeholders who are not involved in the course, 4) *collaborate* with other and their instructor, and 5) engage in *iteration*, or trouble-shooting, problem-solving, or otherwise repeating, revising, or replicating work. For every study, Elicit provided binary (yes/no) output for each feature. We randomly checked 35 studies to verify the Elicit output by examining the full text of the study for evidence for or against presence of each feature in the course design description. We also checked an additional 36 studies that Elicit had marked at least one feature as ‘no’ to verify absence of the feature in the course description. We also examined whether elements of CURE design were measured using the Laboratory Course Assessment Survey (LCAS) (Corwin, Runyon, et al., 2015). The LCAS was designed to measure four of the five features that make CUREs distinct from other experiences (excludes scientific practices), distilling them into three measurable dimensions: opportunities for students to make broadly relevant *discoveries*, engage in *iterative* work, and *collaborate* with peers and instructors. Thus, the LCAS was designed to measure how much courses offer opportunities for students to make broadly relevant discoveries, engage in iterative work, and collaborate with peers and instructors (Corwin et al., 2015). Corwin et al. (2015) provided means and standard deviations for levels of broadly relevant discovery, iteration, and collaboration in a sample of CUREs vs. traditional lab courses. We examined full text of the 135 studies to identify papers that employed LCAS and reported results. When possible, we examined whether the LCAS means reported in our analytic sample fell within one standard deviation of the means reported by Corwin et al., (2015). We summarize our results as counts.

### Comparison or Control Group

To address our second research question (*To what extent do CURE studies include comparison or control groups?*), we reviewed our analytic sample to determine whether each study included a *Comparison or Control Group* in addition to the CURE treatment group, thus making them suitable for meta-analysis. To accomplish this, we reviewed each study to describe the study design, including: (1) the timepoint of data collection (pre/post or post only), thus allowing for assessment of changes due to the intervention; (2) inclusion of comparison or control group(s) (yes, no), thus including a standard for comparison; and (3) whether the study involved randomization of students to conditions (yes, no). We report these results as counts.

### Outcomes

To address our third research question (*What student outcomes are being studied for CUREs, and to what extent are these outcomes being measured and reported in ways that enable meta-analysis?*), we sought to determine which papers in our analytic sample described studies of student outcomes from CUREs. CURE instruction has been linked to a wide range of student outcomes spanning cognitive, behavioral, and affective domains (Krim et al., 2019). Furthermore, these outcomes can be measured using a variety of quantitative and qualitative approaches (e.g., surveys, tests, interviews). Given our goal of conducting a meta-analysis, we focused on studies in which *Outcomes* were measured using quantitative approaches with previously published instruments rather than measures that were developed by the study authors. We identified 48 instruments that were employed in our analytic sample (Table S2). We reviewed each instrument to determine which construct(s) it was designed to measure. We also mapped the constructs onto three larger content domains of (1) attitudes, beliefs, and emotions; (2) experiences; and (3) knowledge, skills, and abilities to provide a big-picture view of the outcomes being studied for CURE instruction.

Two outcomes appeared to have been studied sufficiently in CUREs to enable meta-analysis: science self-efficacy (SE) and science identity (SI). Thus, we sought to determine whether these outcomes had been studied and reported in enough detail to evaluate the effects of CURE instruction. We focused on studies that measured SE and SI using scales reported by (Chemers et al., 2011), (Estrada et al., 2011), or (Hanauer et al., 2016). These scales are versions of each other and thus are likely to be measuring the same constructs (SE and SI) in similar ways, enabling extraction of comparable information for meta-analysis. We cross-referenced this subset of papers to determine the study design (i.e., pre/post measurement and inclusion of a *Comparison or Control Group*) and whether the study reported means, standard deviations (or standard errors), and sample sizes (number of students in the study), which are necessary for calculating the effect of an intervention (CURE instruction) on an outcome (SE or SI). We report these results as counts.

## RESULTS

Here we report the results of our efforts to conduct a meta-analysis aimed at determining the average effects of CURE instruction on student outcomes. We describe our results using the PICO framework to reveal how CUREs are being designed and implemented as an intervention and how CUREs are being studied to determine effects on undergraduate student outcomes.

### CUREs as an intervention

To determine the effect of an intervention on outcome, it is important to clearly and consistently define what constitutes the intervention. The review of our analytic sample (n=135) to describe CUREs as interventions, including their duration, dose, and delivery, indicated considerable variation across the three parameters. We observed a wide variation in the duration of the CUREs being studied and how this information was reported. For example, 111 studies reported durations in semesters, ranging from one to five semesters, and 88 studies reported CURE duration in weeks, with a range of 1 to 75 weeks, and an average of 13.7 ± 9.3 weeks. We also observed a wide variation in the dose of CUREs being studied (n=78 reporting), ranging from 1 to 40 hours per week, with an average of 4.8 ± 4.6 hours (Figure S2).

To describe the CUREs being studied, we relied on the five features originally hypothesized to be distinctive to CURE instruction (Auchincloss et al., 2014). All 135 studies made some reference to at least three of the five features (Figure 2). Specifically, all referenced students’ engagement in one or more scientific practices (Figure 2). Almost all described how the CURE being studied engaged students in work that was relevant beyond the course (n=132) and slightly fewer described how the CURE offered opportunities for students to make discoveries (n=130), despite there being some evidence that relevance and discovery cannot be empirically distinguished (Cooper et al., 2019; Corwin, Runyon, et al., 2015). Most of the studies in our sample also described how students collaborated with each other and their instructor in the CURE (n=134), again despite evidence that current approaches to measure collaboration in course design are not able to distinguish between CURE and non-CURE courses (Beck et al., 2025; Corwin, Runyon, et al., 2015; Zajic et al., 2026). Finally, the CUREs in our sample least often reported opportunities for students to engage in some form of iterative work (n=118), despite at least some evidence that iteration may be more influential than other CURE elements for shaping students’ intentions to continue in science (Corwin et al., 2018).

**Figure 2.**
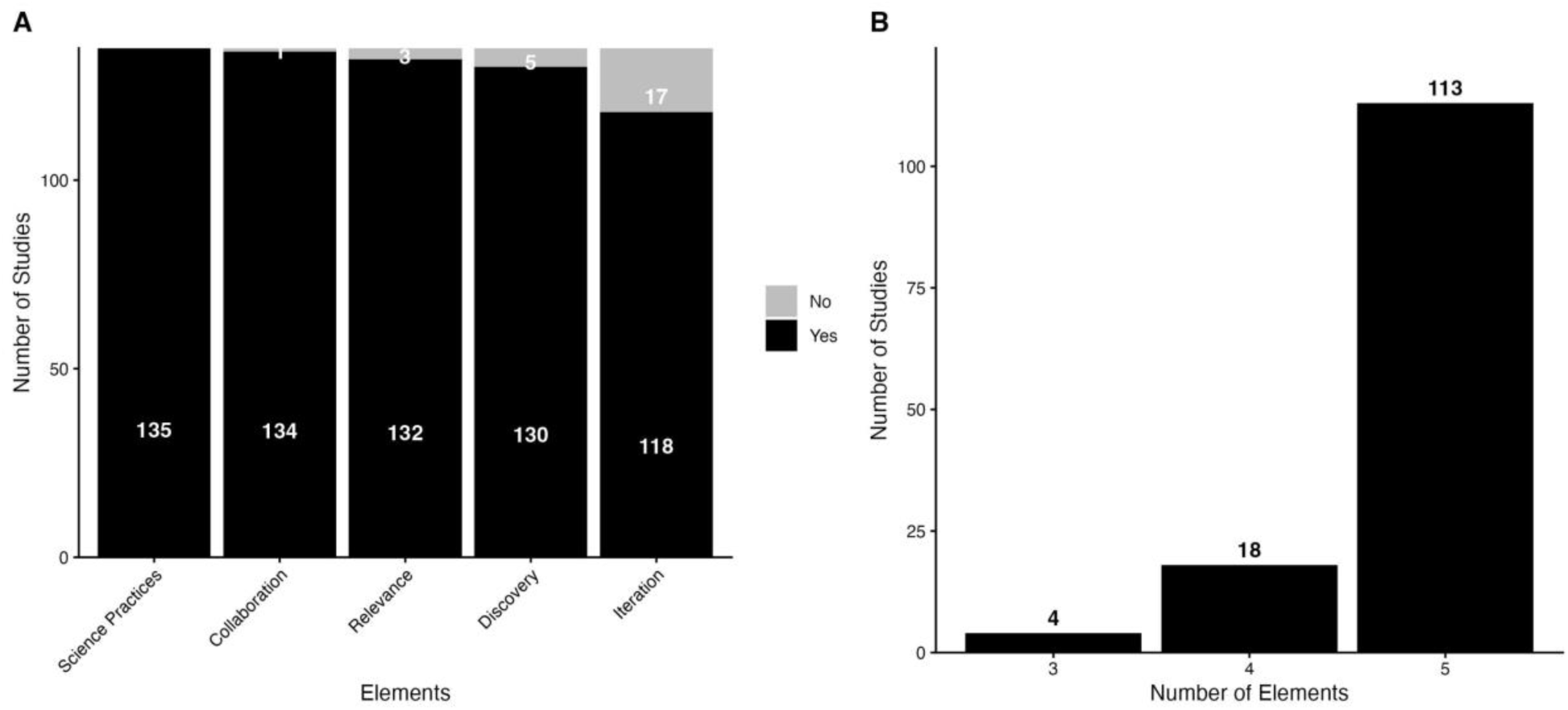
Number of studies in dataset of 135 studies that (A) included each of the five elements hypothesized to make CUREs distinct from other types of instructional experiences (Auchincloss et al., 2014). Numbers within black bars indicate the count of studies that included a given element. (B) Numbers of CURE studies that referenced three, four, or five of the elements.

We noted that only one study examined a design and implementation element beyond those initially proposed by Auchincloss and colleagues (2014). Specifically, Zelaya et al., (2020) examined whether a CURE with an instructor-provided research question (low autonomy) was experienced differently from a CURE in which students chose the research question (high autonomy). Notably, the level of autonomy had no apparent effect on student perceptions of the CURE design, while the duration of the CURE was significantly and positively associated with students’ iterative engagement. Collectively, these results indicate that features of CURE design and implementation largely rely on the framework hypothesized by Auchincloss et al., (2014) but fall short of testing this framework or proposing other elements that may be distinctive to CUREs and influential for students.

Twenty-two studies used the LCAS to measure relevant discovery, iteration, and collaboration. Two of these did not report any LCAS results. Of the 20 that reported results, nine reported means, standard deviations, and sample sizes for *discovery* and *iteration* and 8 reported these values for *collaboration*, permitting some comparison to the published values for CURE instruction from Corwin et al., (2015). Of the nine studies that reported *discovery* scores, eight fell within one standard deviation of the CURE values reported by Corwin et al., (2015). Of the studies reporting *iteration* scores, six studies indicated *iteration* values that fell within one standard deviation values reported by Corwin et al., (2015). We did not examine collaboration scores because Corwin et al. (2015) presented evidence that these scores did not differ between CUREs and non-CUREs. These results suggest that studies of CUREs mostly do not measure or report on the intervention design in ways that facilitate meta-analysis.

Of the 20 studies that reported LCAS results, six included a comparison or control group and reported statistical tests to examine whether LCAS scores from students in CURE differed from students in non-CURE. Four studies found that CURE students reported higher *discovery* than non-CURE students (Callahan et al., 2022; Cooper et al., 2019; D’Arcy et al., 2019; Esparza et al., 2020) and two studies (Allen et al., 2021; Corwin et al., 2018) did not detect differences in reported *discovery* between the groups. The results for *iteration* were also mixed. D’Arcy et al. (2019) and Corwin et al. (2018) observed significantly higher reports of *iteration* for CURE students compared to non-CURE students, while Cooper et al. (2019), and Allen et al. (2021) found no significant differences in *iteration* reported by CURE and non-CURE students. Esparza et al., (2020) and Callahan et al., (2022) did not examine differences in reported *iteration*. These results provide further evidence that CUREs vary in design and implementation and suggest that CUREs are mostly not studied in ways that enable meta-analysis.

### Inclusion of Comparison or Control Groups

To determine the effect of an intervention on outcomes, it is important to compare the intervention to “business as usual” or some other relevant standard (e.g., traditional lab courses, non-CURE lab courses). Of the 135 studies in our analytic sample, about half (n=65) studies included a comparison or control group (Table 1). The comparison or control groups varied widely and included comparisons to published results, proprietary datasets, or results from students in other course types (e.g., traditional lab courses, inquiry lab courses) or experiences (e.g., summer undergraduate research program). These results indicate that studies of CUREs may not be designed in ways that support meta-analysis.

**Table 1.** Study design characteristics of the analytic sample, including data collection timepoints, presence of a comparison group, and randomization status, reported as counts. An additional three studies in single group and three in comparison group were unclear in terms of data collection timepoint.

|  | Single group | Comparison group | Randomized controlled trial |
| --- | --- | --- | --- |
| **Pre/post** | 46 | 36 | 2 |
| **Post only** | 25 | 26 | 0 |

### Measurement and Reporting of Outcomes

Drawing conclusions about the effect(s) of an intervention on outcomes requires a sufficient number of studies relating the CURE to each outcome. The 135 studies in our analytic sample measured a wide range of outcomes (Figure 3) and modest numbers of studies related to each outcome. In an effort to determine whether enough studies of particular outcomes had been conducted for us to conduct a cross-study analysis, we reviewed all 135 studies to determine the target outcome(s) and how it was being measured (i.e., what instrument was used to measure the construct). Figure 3 summarizes the constructs we observed (left), the domains represented by each construct (middle: attitudes, beliefs, and emotions; experiences; knowledge, skills, and abilities), and the instrument used to measure (right). We grouped some constructs to facilitate interpretation of findings. The width of each path from construct to domain to instrument is generally proportional to their frequency in our dataset. The Classroom Undergraduate Research Experience (CURE) Survey, Survey of Undergraduate Research Experiences (SURE), and Research on the Integrated Science Curriculum (RISC) Surveys (Lopatto, 2019) were used in a substantial portion of the studies in our sample. However, we were unable to find information about the target constructs for these instruments. Collectively, these results suggest that CUREs are being taught and studied to shift students’ science-related attitudes, beliefs (e.g., self-efficacy), and/or emotions (e.g., interest) and develop their scientific and academic skills.

**Figure 3:**
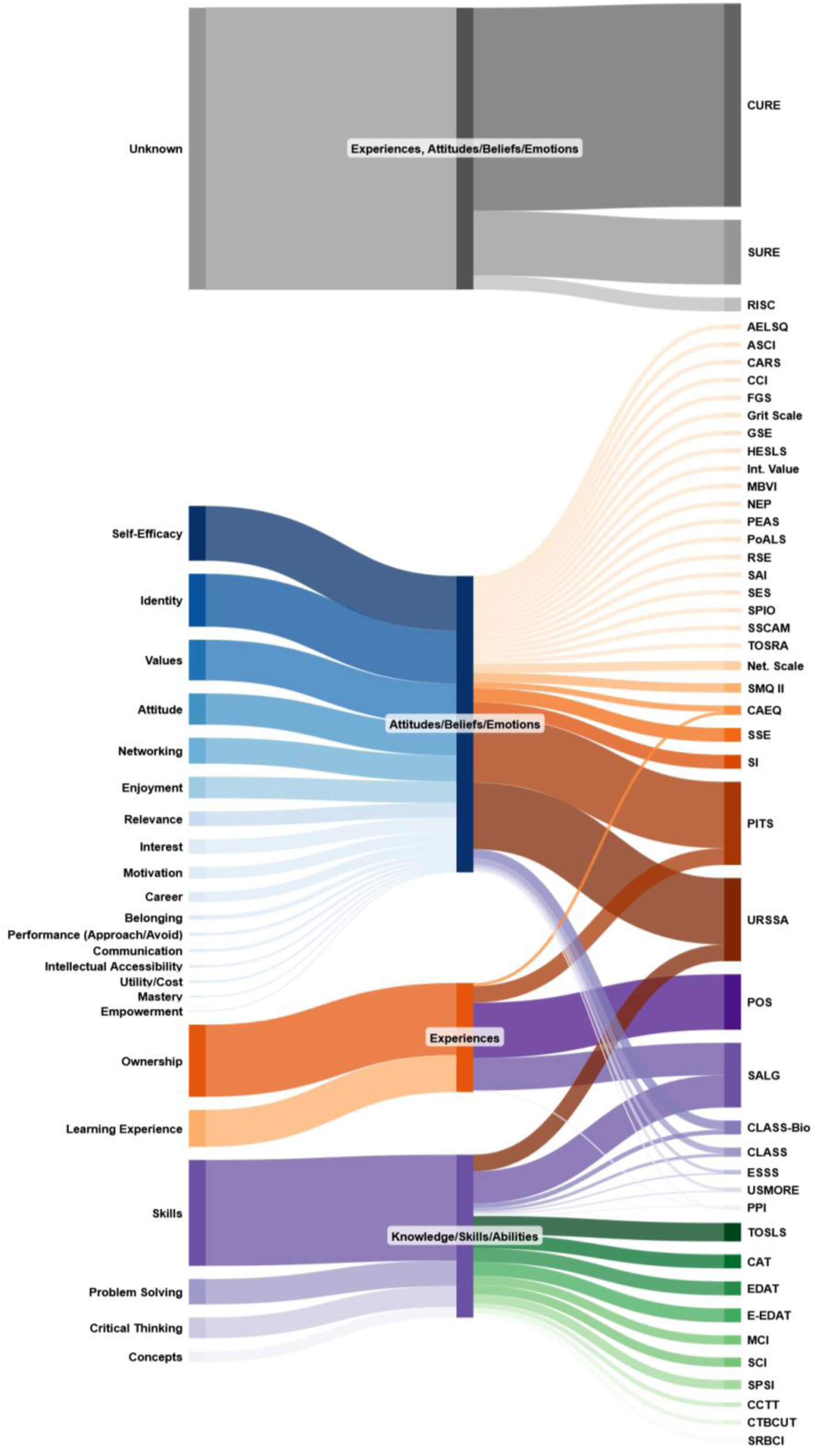
Sankey diagram of published instruments used to study CUREs. Constructs are listed on the left, content domains in the center, and corresponding instruments on the right. Width of the path represents the frequency of a given construct, domain, and instrument within the dataset of 135 studies identified as using a published instrument to measure student outcomes.

To explore whether meta-analysis related to any of these outcomes would be feasible, we reviewed our sample to find the outcomes that were measured and reported most consistently. We identified 18 papers that measured scientific self-efficacy (SE) and/or science identity (SI), citing the persistence in the sciences assessment survey, PITS, (Hanauer et al., 2016) as the instrument, and an additional five papers that measured SE and/or SI, citing (Estrada et al., 2011) or (Chemers et al., 2011). We reviewed these papers more closely and determined that the studies were designed and data were collected and reported in ways that precluded meta-analysis. We observed a range of design, methodology, and reporting issues across the studies, which we describe below.

#### Post only with comparison (between subjects only)

Six studies reported post-course SE and SI for students, referencing some standard for comparison. Two reported post-CURE values for both constructs and did not include either pre-CURE values or another type of course as a standard for comparison. These studies compared their results to published values of SE and SI measured for other CUREs (Johnson et al., 2022) or other lab course types (e.g., Zelaya et al., 2020 compared values from their high and a low autonomy CUREs with published values from the SEA-Phages CURE and traditional lab courses). Four papers reported post-course values of SE and SI for students in a CURE compared with students who completed another type of lab course (Hanauer et al., 2018; Hanauer, Graham, SEA-PHAGES, et al., 2017; LaForge & Martin, 2022; Stanfield et al., 2022). Although these studies reported data (means, standard deviation, sample sizes) that enabled some level of comparison between CURE and non-CURE students, the absence of data on pre-course values in both conditions precluded making causal claims about effects of the course type on students’ SE and SI.

#### Pre/post design with no comparison (within subjects only)

Seven studies compared the SE, SI, or both for students before and after they completed a CURE (Adkins-Jablonsky et al., 2020; Murren et al., 2019; Shuster et al., 2019; Smith et al., 2022; Stovall et al., 2019; Vater et al., 2021; Wilczek et al., 2022). All reported significant gains in SE and SI, and all but one reported their results following best practices for measuring constructs (i.e., at the scale level rather than the item level). Furthermore, all six studies reported results in ways that make future meta-analyses feasible (i.e., inclusion of means, standard deviations, and sample sizes for each variable measured).

#### Pre/post design with comparison (within and between subjects)

Five studies reported students’ SE and/or SI measured before and after a CURE and a comparison experience (Allen et al., 2021; Cole et al., 2021; D’Arcy et al., 2019; Esparza et al., 2020; Ott et al., 2020). While all were designed and reported in ways that could support meta-analysis, each had elements that made it impossible to evaluate the effects of CURE instruction per se. For example, Ott et al. (2020) studied the effects of completing a “research badge” experience coupled with a CURE versus participation in neither. They found that students in the treatment (research badge + CURE) reported significant gains in SE and no difference in SI compared to non-participants, and they reported means, standard deviations, and sample sizes, enabling meta-analysis. Yet, because the treatment included both a CURE and the research badge experience, the specific effect of CURE instruction could not be determined. Cole et al. (2021) compared two different types of CUREs (modular vs. semester-long) rather than comparing changes in SE and SI between students who completed a CURE vs. a non-CURE. Students in both types of CUREs reported similar growth on these constructs. D’Arcy et al. (2019) studied the effect of a CURE on SI compared to a non-CURE course. However, they reported results for only one item on the SI scale rather than treating the scale as a measure of science identity as a latent variable. Finally, Esparza et al. (2020) focused their mixed-methods study on the effects of student and instructor behaviors in CUREs vs. non-CUREs. They investigated how these behaviors related to changes in SE and SI, rather than the effects of course type (CURE vs. non-CURE) on these outcomes. In sum, limitations in study design and methods across this collection of studies precluded meta-analysis.

## DISCUSSION

Our goal at the outset of this study was to conduct a meta-analysis of CURE literature to determine the overall effect of CURE instruction on student outcomes. We identified over 300 CURE publications, 135 of which used a published instrument to study student outcomes. We found substantial limitations in these studies, which precluded meta-analysis. Our results indicate that advancements are needed in the design and reporting of research on CUREs to gauge their effects on student outcomes.

### CUREs are not a singular intervention

To enable meta-analysis of CURE instruction, CUREs must themselves be defined as an educational intervention, including the aim, materials, procedures, duration, frequency, and timing of activities anticipated to impact learners’ outcomes (Harackiewicz & Priniski, 2018). This level of reporting is necessary for two reasons. First, it allows researchers to have confidence that they are studying the intended intervention and attribute outcomes to that intervention. Second, it allows practitioners to replicate the intervention with some hope of achieving similar outcomes. Many of the 135 CURE studies in our dataset fell short of reporting details that would allow for replication or systematic study (Phillips et al., 2016). Phillips and colleagues (2016) proposed that, for an educational intervention to be replicable, authors must provide information on learning objectives, content, delivery mode (e.g., lecture, lab, face-to-face), setting, dose schedule, and duration, along with theoretical underpinnings of the expected outcome(s) (for detailed guidelines and checklists on reporting educational interventions see (Phillips et al., 2016; Upsher et al., 2025)). The intervention description must also include an evaluation of whether the intervention was implemented and experienced as intended (Offerdahl et al., 2018) and produced the expected outcomes (Phillips et al., 2016; Upsher et al., 2025). For the studies in our sample that described CURE instruction in detail, we found substantial variation in their duration, dose, and design, indicating instructors emphasized different aspects of the instructional experience. This result echoes findings from others regarding the wide variation in design and implementation of CURE instruction (Beck et al., 2023; Mendez et al., 2025; Zajic et al., 2026). In essence, the CUREs we reviewed here appear to be distinct interventions.

### Designs of CURE studies limit what can be concluded

Most of the 135 CURE studies in our sample did not include a comparison or control group, and even fewer utilized methods to control for student-level differences across conditions. This is not surprising given the difficulty of designing and executing experimental and quasi-experimental studies in undergraduate education. Yet, to determine the overall effects of CUREs on student outcomes through meta-analysis, CURE studies with comparison or control groups are needed. Ideally, study designs would involve random assignment to condition or other quasi-experimental methods, such as the use of careful matching procedures (e.g., Hanauer et al., 2017; Rodenbusch et al., 2016; Xu & Theobald, 2026). Such designs may be particularly difficult to accomplish with CURE instruction because many institutions are phasing in CUREs to replace all sections of a given course. For example, an institution that aims to ensure all biology majors get at least some research experience may transform all sections of an introductory biology laboratory course into CUREs, leaving no sections that could serve as a standard for comparison. Thus, to more robustly test the effects of CURE instruction on student outcomes, it may be necessary to form cross-institution collaborations to study CURE effectiveness while accounting for student- and institution-level differences. The *Science Study* is an example of how such research could be done; this longitudinal study of the impacts of undergraduate research programming for minoritized students involved careful matching of participating and non-participating campuses and careful testing and accounting for student-level differences (Woodcock et al., 2026). Alternatively, scholars could come to agreement on a set of CURE design and student outcome variables to be measured across lab course contexts and reported in a way that allow for future meta-analysis.

### Measurement methods also limit what can be concluded about CURE effects

Our research revealed two main limitations of how student outcomes of CURE instruction are being studied. First, the studies in our analytic sample explored a wide variety of student outcomes – ranging from critical thinking skills to conceptual knowledge to enjoyment of science. Although each of these outcomes has value, it is unlikely that all CUREs are designed to achieve these outcomes for all participating students. Second, even when studies in our analytic sample presented results related to the same outcome (e.g., scientific self-efficacy), the outcome was measured in a variety of ways that undermined what could be inferred from the data. For example, item wording was sometimes changed without explanation or justification, or items were added to or cut from scales in ways that could affect the meaning of the scale. In addition, few if any analyses were reported to assess whether these changes influenced the validity or reliability of the measurement. To enable future meta-analysis, research on CUREs would benefit from some convergence around outcomes most likely to be realized *across* CUREs rather than within a single CURE. These outcomes should be assessed using instruments with evidence of validity and reliability (Bandalos, 2018; Kane, 1992; McCoach et al., 2013; Tuma & Dolan, 2025) in CURE contexts to enable more defensible conclusions.

### Recommendations for future research

The body of work compiled as a part of this study indicates that there is sustained scholarly interest in studying CUREs, and that CURE instruction has advanced beyond isolated initiatives to an educational approach with broader institutional support. Our results suggest there may now be an opportunity to conduct large-scale investigations *across* institutions to understand the effects of CUREs on student outcomes, including how CURE dose, duration, and design influence student outcomes in diverse contexts. Multi-institution CURE investigations (e.g., Tiny Earth (Hurley et al., 2021), SEA-PHAGES (Hanauer, Graham, SEA-PHAGES, et al., 2017), Bean Beetle CURE (Zelaya et al., 2020), or Squirrel-Net (Connors et al., 2021)) and that include standardized design and data collection would go a long way to generating more generalizable findings. This endeavor could be aided by coalescing on a few theories of change or conceptual models of *how* CUREs influence student outcomes and selecting a *common suite* of instruments to measure the relevant constructs. Prior theoretical work has presented hypotheses that could be leveraged to inform such an effort (Auchincloss et al., 2014; Corwin, Graham, et al., 2015a; Linn et al., 2015).

Addressing the following questions, roughly in order, should be help avoid the methodological shortcomings we observed when studying CURE effects on student outcomes in the future:

– *What is the goal of the CURE?* CUREs may be taught to spark student interest in science or research, to enable students to explore and clarify their career intentions, to broaden student access to research experiences, or to support student persistence and success in science. The goal of the CURE should give both the design and implementation of the CURE itself and the approach and methods used to evaluate whether the goal has been achieved (Cooper et al., 2017).
– *What outcome(s) would indicate the goal has been achieved?* Identifying the affective, behavioral, or cognitive outcomes that provide evidence that the intended goal has been achieved is important for deciding what to measure in a CURE study.
– *What prior research or theory suggests that the intervention could or should lead to the outcomes?* To justify a study, there must be causal logic connecting the intended outcome to the intervention (X aspects of the CURE should cause Y outcomes), and this logic should be grounded in prior theoretical or empirical work. There are various ways to articulate causality, such as by using a theory of change (Reinholz & Andrews, 2020), logic model (Knowlton & Phillips, 2013), or pathway model (Corwin, Graham, et al., 2015b). The key is to formulate a strong rationale supported by prior research of how the intervention is likely to relate to the outcomes *prior* to designing or conducting the study.
– *How can the intervention and outcomes be measured in ways that enable valid and reliable inferences to be made*? Whenever possible, the intervention itself and the intended outcomes should be measured using established approaches with evidence of validity and reliability when used in the intended context with the intended population. Any changes to established measures should be explained and well-justified. It may be useful to collect pilot or preliminary data to ensure measurement tools are operating as intended and useful for detecting changes over time (e.g., no floor or ceiling effects). Timing of data collection should also be considered to ensure the outcomes are being measured at the “right” time.
– *What would be a suitable comparison or control group?* A comparison or control group must be identified to attribute CURE activities to the student outcomes. If random assignment to condition (CURE, non-CURE) is not feasible, it is important to consider what other factors might influence student selection into condition and the outcomes of interest. See Cooper et al. (2019) for an example of how this could be done within a single institution.

### Recommendations for reporting

Meta-analysis of the effects of CUREs on student outcomes will only be possible if reporting of both CURE interventions and student outcomes is improved. We suggest following established standards for reporting educational interventions, such as the CheckList Of Standards of reporting in Education Research (CLOSER), the Checklist for Intervention Description of Education Research (CIDER) (Upsher et al., 2025), or the Guideline for Reporting Evidence-based practice Educational interventions and Teaching (GREET) (Phillips et al., 2016). These checklists are designed to capture enough details for others to fully understand an intervention being studied as well as key aspects of study designs and methods. We also recommend that discipline-based education research journals, which is where the majority of studies in our sample were published, adopt guidelines for reporting quantitative methods and results, such as those outlined by American Psychological Association (Levitt, 2020) or American Educational Research Association (Levine, 2006). Briefly, descriptive statistics (median or mean, range, standard deviation, participant sample size) should be provided for every outcome measured at every timepoint. If instruments measure multiple constructs, descriptive statistics should be provided for each construct.

We observed a preponderance of Likert-type survey data in our analytic sample, and these data were reported in ways that undermined their interpretability and utility. For example, the direction and numerical representation of response options should be labeled what reporting results from these scales (e.g., 1= strongly disagree, 5= strongly agree) (South et al., 2022). One challenge with using scales is that data are inherently ordinal or categorical. Although debates continue about whether scale scores can be treated as continuous, we recommend first visualizing the data to get a sense of whether it is behaving more categorically or continuously. Options for visualization include histograms, violin plots, and stacked bar charts (Koo & Yang, 2025; Sullivan & Artino Jr, 2013). Data should also be evaluated for normality to inform analytic decisions, such as whether to use non-parametric tests (Koo & Yang, 2025; Sullivan & Artino Jr, 2013). Parametric tests may be appropriate, such as when sample sizes are large and data are normally distributed (South et al., 2022; Sullivan & Artino Jr, 2013). Finally, any statistical analysis should be accompanied by a report of the sample size, effect size or odds ratio, estimates of uncertainty, and p-values. Following these guidelines has the potential to improve research on CUREs and position the field for future meta-analysis.

### Limitations

Several limitations should be considered when interpreting the findings described here. First, our search strategy focused on a subset of CURE studies: those that were written in English, that explicitly referred to “course-based undergraduate research” or similar terms, and that were quantitative in nature. As a result, we may have missed studies about the effects of CUREs or similar educational innovations on student outcomes that would not easily lend themselves to meta-analysis. We also limited our in-depth analysis to studies that used a published instrument, but we did not examine how the referenced instrument was deployed in data collection (i.e., whether the instrument was used as intended). This may have influenced our results by excluding informative studies of CUREs that used unpublished instruments or including CURE studies that used instruments in problematic ways. Finally, we used AI as a supportive tool to increase extraction efficiency in reviewing a large body of literature (>300 publications). While Elicit AI states it produces answers with 90% accuracy and we reviewed AI outputs, it is possible that studies were excluded if Elicit did not identify and assessment instrument in the text. Furthermore, as we reviewed the AI output, we identified ambiguity in CURE designs because how authors reported CURE instruction varied considerably from a brief overview of their design to more extensive week-by-week description. To enhance confidence in our results, we shifted to human review once we narrowed our focus to the analytic sample of 135 studies.

## Conclusion

Interest in understanding the effects of CURE instruction is strong as evidenced by the number of studies we identified in the literature. Yet, the variation in how CURE instruction is defined, studied, and reported is so wide that meta-analysis is not feasible and generalizations about CUREs as an intervention are currently not possible. Improvements in study design, methods, and reporting based on the recommendations we made here will go a long way in ensuring that as a community we gather evidence that ultimately allows us to elucidate what aspects of CURE instruction lead to desired outcomes for students.

## Supporting information

Lantz_et_al_Supplemental

## Acknowledgments

This work was supported in part by the Georgia Athletic Association Professorship for Innovative Science Education and by the National Science Foundation Center for Chemical Currencies of a Microbial Planet (OCE-2019589); this is C-CoMP Publication #105. Any opinions, findings, conclusions, or recommendations expressed in this material are those of the authors and do not necessarily reflect the views of any of the funding organizations.

## References

Adkins-Jablonsky, S. J., Akscyn, R., Bennett, B. C., Roberts, Q., & Morris, J. J. (2020). Is community relevance enough? Civic and science identity impact of microbiology CUREs focused on community environmental justice. Frontiers in Microbiology, 11, 578520.

Allen, W. E., Hosbein, K. N., Kennedy, A. M., Whiting, B., & Walker, J. P. (2021). Embedding Research Directly into the Chemistry Curriculum with an Organic to Analytical Sequence. Journal of Chemical Education, 98(7), 2188–2198. 10.1021/acs.jchemed.0c01263

Auchincloss, L. C., Laursen, S. L., Branchaw, J. L., Eagan, K., Graham, M., Hanauer, D. I., Lawrie, G., McLinn, C. M., Pelaez, N., Rowland, S., Towns, M., Trautmann, N. M., Varma-Nelson, P., Weston, T. J., & Dolan, E. L. (2014). Assessment of Course-Based Undergraduate Research Experiences: A Meeting Report. CBE— Life Sciences Education 13(1), 29–40. 10.1187/cbe.14-01-0004

Bandalos, D. L. (2018). Measurement Theory and Applications for the Social Sciences (1 edition). The Guilford Press.

Beck, C. W., Cole, M. F., & Gerardo, N. M. (2023). Can We Quantify If It’s a CURE? Journal of Microbiology & Biology Education. 24:e00210–22. 10.1128/jmbe.00210-22

Beck, C. W., Gerardo, N. M., Karippadath, A., Younge, S. N., & Blumer, L. S. (2025). Effect of Student Autonomy and CURE Duration on Student Perceptions of Research Activities. The American Biology Teacher, 87(6), 334–340.

Buchanan, A. J., & Fisher, G. R. (2022). Current status and implementation of science practices in course-based undergraduate research experiences (CUREs): A systematic literature review. CBE—Life Sciences Education, 21(4), ar83.

Callahan, K. P., Peterson, C. N., Martinez-Vaz, B. M., Huisinga, K. L., Galport, N., Koletar, C., Eddy, R. M., Provost, J. J., Bell, J. K., & Bell, E. (2022). External collaboration results in student learning gains and positive STEM attitudes in CUREs. CBE—Life Sciences Education, 21(4), ar74.

Carson, S. (2015). Targeting critical thinking skills in a first-year undergraduate research course. Journal of Microbiology & Biology Education, 16(2), 148–156.

Chandler, J., Cumpston, M., Li, T., Page, M. J., & Welch, V. (2019). Cochrane handbook for systematic reviews of interventions. Hoboken: Wiley, 4(1002), 14651858.

Chemers, M. M., Zurbriggen, E. L., Syed, M., Goza, B. K., & Bearman, S. (2011). The Role of Efficacy and Identity in Science Career Commitment Among Underrepresented Minority Students. Journal of Social Issues, 67(3), 469–491. 10.1111/j.1540-4560.2011.01710.x

Cole, M. F., Hickman, M. A., & Morran, L. (2021). Assessment of course-based research modules based on faculty research in introductory biology (Pt. E0014821). Journal of Microbiology & Biology Education, 22(2) 10–1128. 10.1128/jmbe.00148-21

Connors, P. K., Lanier, H. C., Erb, L. P., Varner, J., Dizney, L., Flaherty, E. A., Duggan, J. M., Yahnke, C. J., & Hanson, J. D. (2021). Connected while distant: Networking CUREs across classrooms to create community and empower students. Integrative and Comparative Biology, 61(3), 934–943.

Cooke, A., Smith, D. M., & Booth, A. (2012). The benefits of a systematic search strategy when conducting qualitative evidence synthesis; the SPIDER tool. Qualitative Health Research, 22(10), 1435–1443.

Cooper, K. M., Blattman, J. N., Hendrix, T., & Brownell, S. E. (2019). The Impact of Broadly Relevant Novel Discoveries on Student Project Ownership in a Traditional Lab Course Turned CURE. CBE—Life Sciences Education, 18(4), ar57. 10.1187/cbe.19-06-0113

Cooper, K. M., Soneral, P. A. G., & Brownell, S. E. (2017). Define Your Goals Before You Design a CURE: A Call to Use Backward Design in Planning Course-Based Undergraduate Research Experiences. Journal of Microbiology & Biology Education, 18(2). 10.1128/jmbe.v18i2.1287

Corwin, L. A., Graham, M. J., & Dolan, E. L. (2015a). Modeling course-based undergraduate research experiences: An agenda for future research and evaluation. CBE—Life Sciences Education, 14(1), es1.

Corwin, L. A., Graham, M. J., & Dolan, E. L. (2015b). Modeling Course-Based Undergraduate Research Experiences: An Agenda for Future Research and Evaluation. CBE-Life Sciences Education, 14(1), es1. 10.1187/cbe.14-10-0167

Corwin, L. A., Runyon, C. R., Ghanem, E., Sandy, M., Clark, G., Palmer, G. C., Reichler, S., Rodenbusch, S. E., & Dolan, E. L. (2018). Effects of Discovery, Iteration, and Collaboration in Laboratory Courses on Undergraduates’ Research Career Intentions Fully Mediated by Student Ownership. CBE—Life Sciences Education, 17(2). 10.1187/cbe.17-07-0141

Corwin, L. A., Runyon, C., Robinson, A., & Dolan, E. L. (2015). The laboratory course assessment survey: A tool to measure three dimensions of research-course design. CBE—Life Sciences Education, 14(4), ar37.

D’Arcy, C. E., Martinez, A., Khan, A. M., & Olimpo, J. T. (2019). Cognitive and Non-Cognitive Outcomes Associated with Student Engagement in a Novel Brain Chemoarchitecture Mapping Course-Based Undergraduate Research Experience. Journal of Undergraduate Neuroscience Education, 18(1), A15–a43.

DeChenne-Peters, S. E., Rakus, J. F., Parente, A. D., Mans, T. L., Eddy, R., Galport, N., Koletar, C., Provost, J. J., Bell, J. E., & Bell, J. K. (2023). Length of course-based undergraduate research experiences (CURE) impacts student learning and attitudinal outcomes: A study of the Malate dehydrogenase CUREs Community (MCC). Plos One, 18(3), e0282170.

Esparza, D., Wagler, A. E., & Olimpo, J. T. (2020). Characterization of instructor and student behaviors in CURE and non-CURE learning environments: Impacts on student motivation, science identity development, and perceptions of the laboratory experience. CBE—Life Sciences Education, 19(1), ar10.

Estrada, M., Woodcock, A., Hernandez, P. R., & Schultz, P. (2011). Toward a model of social influence that explains minority student integration into the scientific community. Journal of Educational Psychology, 103(1), 206.

Goodwin, E. C., Anokhin, V., Gray, M. J., Zajic, D. E., Podrabsky, J. E., & Shortlidge, E. E. (2021). Is This Science? Students’ Experiences of Failure Make a Research-Based Course Feel Authentic. CBE-Life Science Education, 20(1). 10.1187/cbe.20-07-0149

Hanauer, D. I., Graham, M. J., SEA-PHAGES, Betancur, L., Bobrownicki, A., Cresawn, S. G., Garlena, R. A., Jacobs-Sera, D., Kaufmann, N., Pope, W. H., Russell, D. A., Jacobs, W. R., Sivanathan, V., Asai, D. J., Hatfull, G. F., Actis, L., Adair, T., Adams, S., Alvey, R., … Zimmerman, A. (2017). An inclusive Research Education Community (iREC): Impact of the SEA-PHAGES program on research outcomes and student learning. Proceedings of the National Academy of Sciences, 114(51), 13531–13536. 10.1073/pnas.1718188115

Hanauer, D. I., Graham, M. J., Sea-Phages, Betancur, L., Bobrownicki, A., Cresawn, S. G., Garlena, R. A., Jacobs-Sera, D., Kaufmann, N., Pope, W. H., Russell, D. A., Jacobs, W. R., Sivanathan, V., Asai, D. J., Hatfull, G. F., Actis, L., Adair, T., Adams, S., Alvey, R., … Zimmerman, A. (2017). An inclusive research education community (iREC): Impact of the SEA-PHAGES program on research outcomes and student learning. Proceedings of the National Academy of Sciences, 114(51), 13531–13536. 10.1073/pnas.1718188115

Hanauer, D. I., Nicholes, J., Liao, F.-Y., Beasley, A., & Henter, H. (2018). Short-term research experience (SRE) in the traditional lab: Qualitative and quantitative data on outcomes. CBE—Life Sciences Education, 17(4), ar64.

Hanauer, David I., Graham, Mark J., Hatfull, Graham F., & Brickman, Peggy. (2016). A Measure of College Student Persistence in the Sciences (PITS). CBE—Life Sciences Education, 15(4), ar54. 10.1187/cbe.15-09-0185

Harackiewicz, J. M., & Priniski, S. J. (2018). Improving student outcomes in higher education: The science of targeted intervention. Annual Review of Psychology, 69, 409–435.

Hurley, A., Chevrette, M. G., Acharya, D. D., & Lozano, G. L. (2021). Tiny earth: A big idea for stem education and antibiotic discovery. MBio, 12(1), e03432–20. 10.1128/mbio.03432-20

Johnson, K. C., Sabel, J. L., Cole, J., Pruett, C. L., Plymale, R., & Reyna, N. S. (2022). From genetics to biotechnology: Synthetic biology as a flexible course-embedded research experience. Biochemistry and Molecular Biology Education, 50(6), 580–591. 10.1002/bmb.21662

Kane, M. T. (1992). An argument-based approach to validity. Psychological Bulletin, 112(3), 527–535. 10.1037/0033-2909.112.3.527

Knowlton, L. W., & Phillips, C. C. (2013). The logic model guidebook: Better strategies for great results. Sage.

Koo, M., & Yang, S.-W. (2025). Likert-Type Scale. Encyclopedia, 5(1), 18. 10.3390/encyclopedia5010018

Krim, J. S., Coté, L. E., Schwartz, R. S., Stone, E. M., Cleeves, J. J., Barry, K. J., Burgess, W., Buxner, S. R., Gerton, J. M., Horvath, L., Keller, J. M., Lee, S. C., Locke, S. M., & Rebar, B. M. (2019). Models and Impacts of Science Research Experiences: A Review of the Literature of CUREs, UREs, and TREs. CBE—Life Sciences Education, 18(4), ar65. 10.1187/cbe.19-03-0069

LaForge, J., & Martin, E. C. (2022). Impact of Authentic Course-Based Undergraduate Research Experiences (CUREs) On Student Understanding in Introductory Biology Laboratory Courses. American Biology Teacher, 84(3), 137–142. 10.1525/abt.2022.84.3.137

Levine, F. (2006). Standards for reporting on empirical social science research in AERA publications. Educational Researcher, 35, 33–40.

Levitt, H. M. (2020). Reporting qualitative research in psychology: How to meet APA style journal article reporting standards. American Psychological Association.

Linn, M. C., Palmer, E., Baranger, A., Gerard, E., & Stone, E. (2015). Undergraduate research experiences: Impacts and opportunities. Science, 347(6222), 1261757. 10.1126/science.1261757

Lopatto, D. (2019). Undergraduate Research Experience Surveys. https://sure.sites.grinnell.edu/

Maynard, R. (2024). Improving the usefulness and use of meta-analysis to inform policy and practice. Evaluation Review, 48(3), 515–543.

McCoach, D. B., Gable, R. K., & Madura, J. P. (2013). Instrument development in the affective domain. *New York, NY*: *Springer*. *Doi*, *10*, 978–1.

Mendez, Y. C. M., Listyg, B. S., Mardones-Segovia, C., Bowers, B., Cotto, K. M. C., Hilton, L., Koscik, I., Outlaw, B., Portner, S., & Barekzi, N. (2025). Unveiling Undergraduate Research: Employing Ecological Momentary Assessment to Characterize and Compare Undergraduate Research Experiences. CBE—Life Sciences Education, 24(4), ar49.

Merkle, J. A., Devergne, O., Kelly, S. M., & Croonquist, P. A. (2023). Fly-CURE, a Multi-institutional CURE using Drosophila, Increases Students’ Confidence, Sense of Belonging, and Persistence in Research. bioRxiv. 10.1101/2023.01.16.524319.abstract

Methley, A. M., Campbell, S., Chew-Graham, C., McNally, R., & Cheraghi-Sohi, S. (2014). PICO, PICOS and SPIDER: a comparison study of specificity and sensitivity in three search tools for qualitative systematic reviews. BMC Health Services Research, 14(1), 579.

Murren, C. J., Wolyniak, M. J., Rutter, M. T., Bisner, A. M., Callahan, H. S., Strand, A. E., & Corwin, L. A. (2019). Undergraduates Phenotyping Arabidopsis Knockouts in a Course-Based Undergraduate Research Experience: Exploring Plant Fitness and Vigor Using Quantitative Phenotyping Methods. Journal of Microbiology & Biology Education, 20(2). 10.1128/jmbe.v20i2.1650

National Academies of Sciences, Engineering, and Medicine. (2015). Integrating discovery-based research into the undergraduate curriculum: Report of a convocation. Washington, DC: National Academies Press.

Offerdahl, E. G., McConnell, M., & Boyer, J. (2018). Can I have your recipe? Using a fidelity of implementation (FOI) framework to identify the key ingredients of formative assessment for learning. CBE—Life Sciences Education, 17(4), es16.

Olimpo, J. T., Fisher, G. R., & DeChenne-Peters, S. E. (2016). Development and Evaluation of the Tigriopus Course-Based Undergraduate Research Experience: Impacts on Students’ Content Knowledge, Attitudes, and Motivation in a Majors Introductory Biology Course. CBE-Life Science Education, 15(4). 10.1187/cbe.15-11-0228

Ott, L. E., Godsay, S., Stolle-McAllister, K., Kowalewski, C., Maton, K. I., & LaCourse, W. R. (2020). Introduction to research: A scalable, online badge implemented in conjunction with a classroom-based undergraduate research experience (CURE) that promotes students matriculation into mentored undergraduate research. UI Journal, 11(1), 1–25.

Page, M. J., McKenzie, J. E., Bossuyt, P. M., Boutron, I., Hoffmann, T. C., Mulrow, C. D., Shamseer, L., Tetzlaff, J. M., Akl, E. A., & Brennan, S. E. (2021). The PRISMA 2020 statement: An updated guideline for reporting systematic reviews. Bmj, 372.

Pellegrini, M., Day, E., Scarbrough, H. F., & Pigott, T. D. (2025). A meta-review of education meta-analyses: Relevance, applicability, and accessibility of findings. AERA Open, 11, 23328584251389562.

Phillips, A. C., Lewis, L. K., McEvoy, M. P., Galipeau, J., Glasziou, P., Moher, D., Tilson, J. K., & Williams, M. T. (2016). Development and validation of the guideline for reporting evidence-based practice educational interventions and teaching (GREET). BMC Medical Education, 16(1), 237. 10.1186/s12909-016-0759-1

Pigott, T. D., & Polanin, J. R. (2020). Methodological guidance paper: High-quality meta-analysis in a systematic review. Review of Educational Research, 90(1), 24–46.

R Core Team. (2026). R: A Language and Environment for Statistical Computing [Computer software].

Reinholz, D. L., & Andrews, T. C. (2020). Change theory and theory of change: What’s the difference anyway? International Journal of STEM Education, 7(1), 2. 10.1186/s40594-020-0202-3

Richardson, W. S., Wilson, M. C., Nishikawa, J., & Hayward, R. S. (1995). The well-built clinical question: A key to evidence-based decisions. ACP Journal Club, 123(3), A12–3.

Rodenbusch, S. E., Hernandez, P. R., Simmons, S. L., & Dolan, E. L. (2016). Early Engagement in Course-Based Research Increases Graduation Rates and Completion of Science, Engineering, and Mathematics Degrees. CBE-Life Science Education, 15(2). 10.1187/cbe.16-03-0117

Russell, C. B., & Weaver, G. (2008). Student perceptions of the purpose and function of the laboratory in science: A grounded theory study. International Journal for the Scholarship of Teaching and Learning, 2(2), 9.

Shaffer, C. D., Alvarez, C., Bailey, C., Barnard, D., Bhalla, S., Chandrasekaran, C., Chandrasekaran, V., Chung, H.-M., Dorer, D. R., & Du, C. (2010). The genomics education partnership: Successful integration of research into laboratory classes at a diverse group of undergraduate institutions. CBE—Life Sciences Education, 9(1), 55–69.

Shuster, M. I., Curtiss, J., Wright, T. F., Champion, C., Sharifi, M., & Bosland, J. (2019). Implementing and evaluating a course-based undergraduate research experience (CURE) at a Hispanic-serving institution. Interdisciplinary Journal of Problem-Based Learning, 13(2).

Smith, M. A., Olimpo, J. T., & Santillan, K. A. (2022). Addressing foodborne illness in Côte d’Ivoire: Connecting the classroom to the community through a nonmajors course-based undergraduate research experience. Journal of Microbiology & Biology Education, 23(1), e00212–21. 10.1128/jmbe.00212-21

South, L., Saffo, D., Vitek, O., Dunne, C., & Borkin, M. A. (2022). Effective use of Likert scales in visualization evaluations: A systematic review. 41(3), 43–55.

Stanfield, E., Slown, C. D., Sedlacek, Q., & Worcester, S. E. (2022). A Course-Based Undergraduate Research Experience (CURE) in Biology: Developing Systems Thinking through Field Experiences in Restoration Ecology. CBE-Life Sciences Education, 21(2). 10.1187/cbe.20-12-0300

Stovall, G. M., Huynh, V., Engelman, S., & Ellington, A. D. (2019). Aptamers in education: Undergraduates make aptamers and acquire 21st century skills along the way. Sensors, 19(15), 3270.

Sullivan, G. M., & Artino Jr, A. R. (2013). Analyzing and interpreting data from Likert-type scales. Journal of Graduate Medical Education, 5(4), 541–542.

Treibergs, K. A., Stetzer, M. R., Olson, A. N., Schmid, K., Adjei-Opong, T., Onimode, R., Noyes, K., Eldermire, E. R. B., Couch, B. A., & Smith, M. K. (2025). A Scoping Review of Published Lesson Plans Showcases Two Decades of Change in Undergraduate Life Science Education Resources. CBE—Life Sciences Education, 24(4), ar40. 10.1187/cbe.25-04-0068

Tuma, T. T., & Dolan, E. L. (2025). So, You Want to Measure Something? An Introduction to Measurement Validity in Educational Research. Scholarship and Practice of Undergraduate Research, 9(1), 18–24.

Upsher, R., Dommett, E., Carlisle, S., Conner, S., Codina, G., Nobili, A., & Byrom, N. C. (2025). Improving reporting standards in quantitative educational intervention research: Introducing the CLOSER and CIDER checklists. Journal of New Approaches in Educational Research, 14(1), 2. 10.1007/s44322-024-00022-9

Vater, A., Mayoral, J., Nunez-Castilla, J., Labonte, J. W., Briggs, L. A., Gray, J. J., Makarevitch, I., Rumjahn, S. M., & Siegel, J. B. (2021). Development of a Broadly Accessible, Computationally Guided Biochemistry Course-Based Undergraduate Research Experience. Journal of Chemical Education, 98(2), 400–409. 10.1021/acs.jchemed.0c01073

Watts, F. M., & Rodriguez, J.-M. G. (2023). A Review of Course-Based Undergraduate Research Experiences in Chemistry. Journal of Chemical Education, 100(9), 3261–3275. 10.1021/acs.jchemed.3c00570

Wilczek, L. A., Clarke, A. J., Martinez, M. D. G., & Morin, J. B. (2022). Catalyzing the Development of Self-Efficacy and Science Identity: A Green Organic Chemistry CURE. Journal of Chemical Education, 99(12), 3878–3887. 10.1021/acs.jchemed.2c00352

Woodcock, A., Aguilar, S. D., Hernandez, P. R., Peterson, M. E., & Schultz, P. W. (2026). Broadening participation: 20-year outcomes from undergraduate science training programs. Science Advances, 12(25), eaeh0739.

Xu, S., & Theobald, E. J. (2026). Toward Causal Inferences in Discipline-Based Education Research: Using Regression Discontinuity Design to Understand the Effect of Classroom Interventions. CBE—Life Sciences Education, 25(1), rm1.

Zajic, C. J., Subramanian, K., Adulla, A., Allen, Z., Blitchington, M. B., Brotzman, A., Carrillo, E., Dhruv, J., Evans, T., & Haider, S. (2026). Could instructor talk drive CURE effectiveness? A comparative study of instructor talk in introductory lab courses. CBE—Life Sciences Education, 25(1), ar3.

Zelaya, A. J., Gerardo, N. M., Blumer, L. S., & Beck, C. W. (2020). The Bean Beetle Microbiome Project: A Course-Based Undergraduate Research Experience in Microbiology. Frontiers in Microbiology, 11. 10.3389/fmicb.2020.577621

