## Supplementary material for "Variation in CURE instruction and limitations of CURE studies undermine what can be concluded about CURE effects on student outcomes": Lantz_et_al_Supplemental

**Table S1.** Variables extracted from studies along with instructions and answer structure provided to Elicit AI tool. Example outputs for ‘any answer’ structure.

| **Variable** | **Instructions** | **Answer Structure** | **Example output from Elicit** |
| --- | --- | --- | --- |
| Assessment instrument | Analyze text and identify the type of quantitative data collected on the students. Specify method used (e.g., likert scale surveys, tests, quizzes). Do not include information on the data collected by the students. If quantitative data is not mentioned, leave the answer blank | Any answer | *“Likert scale surveys (Undergraduate Research Student Self-Assessment - URSSA)”* |
| [*48 instruments identified from previous variable outputs*]  Example: PITS | Analyze text and indicate if "Persistence in the Sciences", or "PITS" was mentioned in the study. It has to be explicitly mentioned to indicate 'yes'. | Yes, No |  |
| Use of science practices | Analyze the text and indicate if the research that the students conduct for the CURE includes "use of science practices". Indicate “yes” if there are scientific methods (data collection, experiment design, hypothesis, etc.). Here are similar terms: “the nature of science” and “process of research”. | Yes, No |  |
| Iteration | Analyze the text and indicate if the research that the students conduct for the CURE includes "iteration". | Yes, No |  |
| Discovery | Analyze the text and indicate if the research that the students conduct for the CURE includes "discovery", or that it addresses questions where the answer is unknown within the broader scientific community. | Yes, No |  |
| Relevance | Analyze the text and indicate if the research that the students conduct for the CURE includes "relevance", or that it addresses questions where the answer is unknown within the broader scientific community. | Yes, No |  |
| Collaboration | Analyze the text and indicate if the research that the students conduct for the CURE includes "collaboration", or that students work in teams or small groups on the project. | Yes, No |  |
| Duration | Analyze the text and provide the duration of the course described in the study. Provide number of semesters and number of weeks in the following format  semesters:  weeks:  If the course repeats, give the duration of a single course. | Any answer | *“semesters: 1  weeks: 15”* |
| Hours | Analyze the text and provide the duration of the course in hours per week. For example, a course might meet once a week for 3 hours. | Any answer | *“Not mentioned (the paper does not provide information on the duration of the course in hours per week)”* |

**Table S2.** Assessment instruments implemented in the 135 CURE studies included in our dataset, along with the acronym (when provided) and count of studies using each instrument.

| **Instrument** | **Acronym** | **Number of studies reporting** |
| --- | --- | --- |
| Classroom Undergraduate Research Experience | CURE | 44 |
| Persistence in the Sciences | PITS | 18 |
| Undergraduate Student Self-Assessment Instrument | URSSA | 18 |
| Student Assessment of Learning Gains | SALG | 14 |
| Survey of Undergraduate Research Experiences (I, II or III) | SURE | 14 |
| Project Ownership Survey | POS | 12 |
| Test of Scientific Literacy Skills | TOSLS | 4 |
| Critical Thinking Assessment Test | CAT | 3 |
| Colorado Learning Attitudes about Science Survey | CLASS-bio | 3 |
| Experimental Design Ability Test | E-DAT | 3 |
| Expanded Experimental Design Ability Test | E-EDAT | 3 |
| Research on the Integrated Science Curriculum | RISC | 3 |
| Science identity |  | 3 |
| Science self-efficacy |  | 3 |
| Science Motivation Questionnaire II | SMQ-II | 3 |
| Chemistry Attitudes and Experiences Questionnaire | CAEQ | 2 |
| Colorado Learning Attitudes about Science Survey | CLASS | 2 |
| Microbiology Concept Inventory | MCI | 2 |
| Networking Scale |  | 2 |
| Skills and Concepts Inventory |  | 2 |
| Affective Elements of Science Learning Questionnaire | AESLQ | 1 |
| Attitude toward the Study of Chemistry Inventory | ASCI | 1 |
| Civic Attitudes about the Relevance of Science | CARS | 1 |
| Cornell Critical Thinking Test | CCTT | 1 |
| Classroom Community Inventory |  | 1 |
| Critical Thinking Basic Concepts & Understanding Test |  | 1 |
| Employable Self-Efficacy Survey |  | 1 |
| Future Goals Survey |  | 1 |
| Grit Scale |  | 1 |
| General Self-Efficacy | GSE | 1 |
| Higher Ed Service-Learning Survey | HESLS | 1 |
| Interest value |  | 1 |
| Math-Biology Values Instrument |  | 1 |
| New Ecological Paradigm Scale |  | 1 |
| Participant Perception Indicator | PPI | 1 |
| Patterns of Adaptive Learning Scales | PALS | 1 |
| Pittsburgh Engineering Attitudes Scale-Revised |  | 1 |
| Research Self-Efficacy Scale |  | 1 |
| Science Process Skills Inventory |  | 1 |
| Scientific Attitudes Inventory |  | 1 |
| Self-efficacy Survey | SES | 1 |
| Student Subjective Science Attitude Change Measure | SSACM | 1 |
| Statistical Reasoning in Biology Concepts Inventory |  | 1 |
| STEM Professional Identity Overlap measure |  | 1 |
| Test of Science-Related Attitudes | TOSRA | 1 |
| Undergraduate Scientists-Measuring Outcomes of Research Experiences | USMORE | 1 |

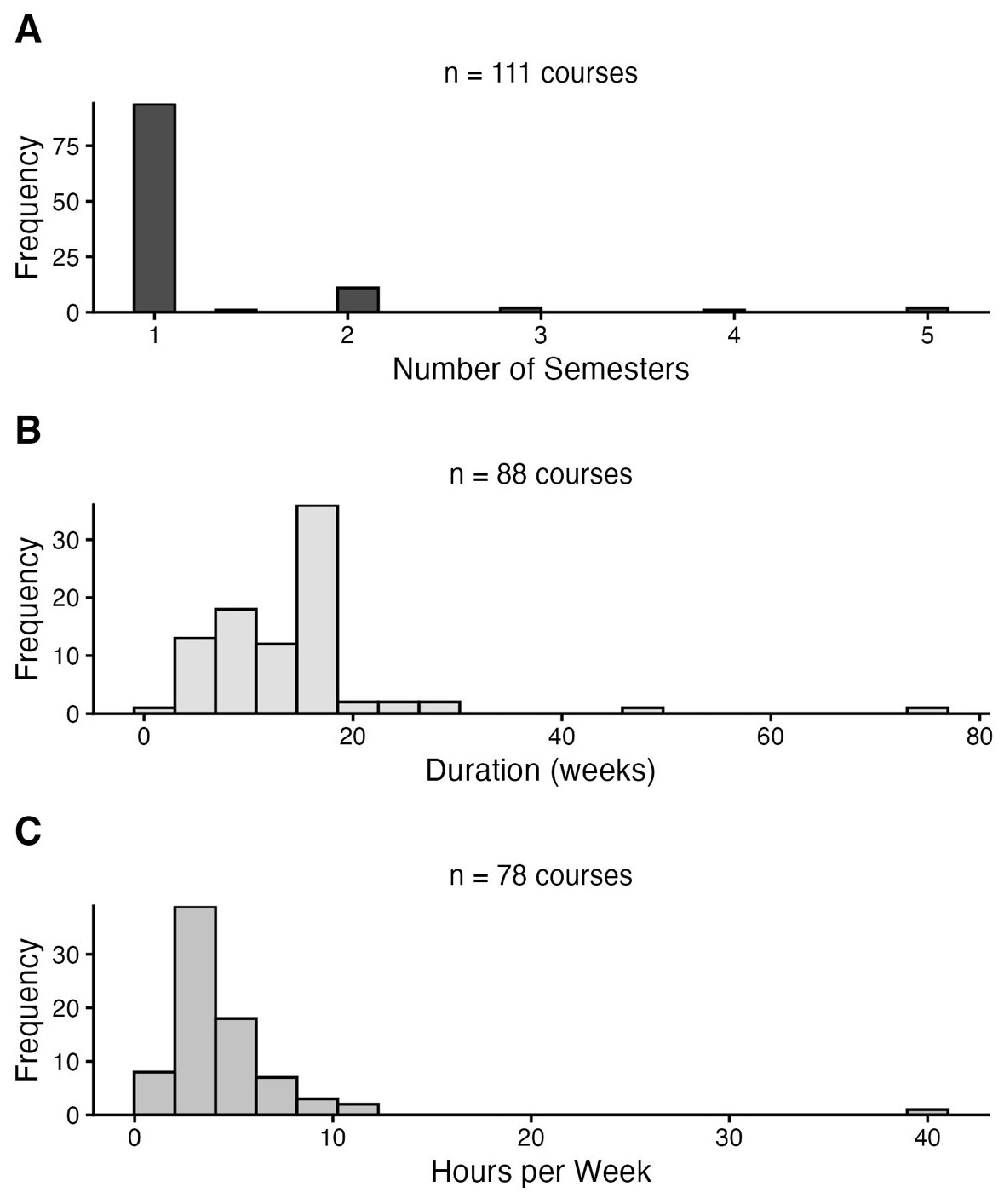

**Figure S1.** Histograms of reported CURE durations in (A) semesters, (B) weeks and (C) hours per week.

**Reference list for the 135 studies included in the dataset.**

Adkins-Jablonsky SJ, Akscyn R, Bennett BC, Roberts Q, Morris JJ. 2020. Is community relevance enough? Civic and science identity impact of microbiology cures focused on community environmental justice. Frontiers in Microbiology. 11:578520.

Al-Ghadhban S, Muqaibel A, Alregib G, Al-Shaikhi A. 2018. Seeding undergraduate research experience: From georgia tech to kfupm case study. International Journal of Electrical Engineering Education. 55(4):313–323.

Allen J, Kuehn S, Creamer E. 2020a. A boost for the cure: Improving learning outcomes with curriculum-based undergraduate research. GSA Today. 30:28–29.

Allen JL, Kuehn SC, Creamer EC, Austin JE. 2020b. A multisemester, curriculum-embedded undergraduate research experience (ms-cure) in the geosciences: Contextual factors influencing student motivation for pursuing research. SPUR-Scholarship and Practice of Undergraduate Research. 4(2):35–43.

Allen WE, Hosbein KN, Kennedy AM, Whiting B, Walker JP. 2021a. Design and implementation of an organic to analytical cure sequence. Journal of Chemical Education. 98(7):2199–2208.

Allen WE, Hosbein KN, Kennedy AM, Whiting B, Walker JP. 2021b. Embedding research directly into the chemistry curriculum with an organic to analytical sequence. Journal of Chemical Education. 98(7):2188–2198.

Arnold DM, Mortensen CJ, Thoron AC, Miller-Cushon EK, Miot JK. 2019. Contrasting science learning gains and attitudes of students in an early research-based experience. NACTA Journal. 63(2):180–187.

Arnold DM, Mortensen CJ, Thoron AC, Miot JK, Miller-Cushon EK. 2018. Identifying the optimal course delivery platform in an undergraduate animal behavior research course. Translational Animal Science. 2(3):311–318.

Ayella A, Beck MR. 2018. A course-based undergraduate research experience investigating the consequences of nonconserved mutations in lactate dehydrogenase. Biochemistry and molecular biology education. 46(3):285–296.

Ballen CJ, Thompson SK, Blum JE, Newstrom NP, Cotner S. 2018. Discovery and broad relevance may be insignificant components of course-based undergraduate research experiences [cures] for non-biology majors. Journal of Microbiology & Biology Education. 19(2).

Bangera G, Harrington K, Shaver I. 2022. Hands-on, hands-off: The community college genomics (comgen) course-based undergraduate research experience. CourseSource. 9.

Bergstrom RA. 2018. Building a new translational research program with undergraduates: A student-driven research class. Science education and civic engagement. 10(1).

Bixby TJ, Miliauskas MM. 2022. Assessment of the short-term outcomes of a semester-long cure in general chemistry lab. Journal of Chemical Education. 99(12):3849–3857.

Bucklin CJ, Mauger L. 2022. Cures: How to create & incorporate a collaborative ant-based project to teach science practices. American Biology Teacher. 84(6):353–357.

Burnette JM, Wessler SR. 2013. Transposing from the laboratory to the classroom to generate authentic research experiences for undergraduates. Genetics. 193(2):367.

Chase AM, Clancy HA, Lachance RP, Mathison BM, Chiu MM, Weaver GC. 2017. Improving critical thinking via authenticity: The caspie research experience in a military academy chemistry course. Chemistry Education Research and Practice. 18(1):55–63.

Clyne AM, Shieh AC, Stanford JS. 2019. A course-based undergraduate research experience in biofluid mechanics. Journal of Biomechanical Engineering-Transactions of the ASME. 141(12).

Cole MF, Hickman MA, Morran L, Beck CW. 2021. Assessment of course-based research modules based on faculty research in introductory biology. Journal of Microbiology & Biology Education. 22(2):10–1128.

Cookmeyer DL, Winesett ES, Kokona B, Huff AR, Aliev S, Bloch NB, Bulos JA, Evans IL, Fagre CR, Godbe KN et al. 2017. Uncovering protein-protein interactions through a team-based undergraduate biochemistry course. Plos Biology. 15(11).

Cooper KM, Blattman JN, Hendrix T, Brownell SE. 2019. The impact of broadly relevant novel discoveries on student project ownership in a traditional lab course turned cure. CBE-Life Sciences Education. 18(4):ar57.

Cooper KM, Knope ML, Munstermann MJ, Brownell SE. 2020. Students who analyze their own data in a course-based undergraduate research experience (cure) show gains in scientific identity and emotional ownership of research. Journal of Microbiology & Biology Education. 21(3).

Corwin LA, Runyon CR, Ghanem E, Sandy M, Clark G, Palmer GC, Reichler S, Rodenbusch SE, Dolan EL. 2018. Effects of discovery, iteration, and collaboration in laboratory courses on undergraduates' research career intentions fully mediated by student ownership. CBE-Life Sciences Education. 17(2):ar20.

Coticone S, Houten L. 2020. Integrating course-based undergraduate research experiences (cures) in advanced forensic science curriculum as an active learning strategy. The Journal of Forensic Science Education 2(2).

Cruz CL, Holmberg-Douglas N, Onuska NPR, McManus JB, MacKenzie IA, Hutson BL, Eskew NA, Nicewicz DA. 2020. Development of a large-enrollment course-based research experience in an undergraduate organic chemistry laboratory: Structure-function relationships in pyrylium photoredox catalysts. Journal of Chemical Education. 97(6):1572–1578.

D'Arcy CE, Martinez A, Khan AM, Olimpo JT. 2019. Cognitive and non-cognitive outcomes associated with student engagement in a novel brain chemoarchitecture mapping course-based undergraduate. Journal of Undergraduate Neuroscience Education. 18(1):A15–A43.

Dahlberg CL, Wiggins BL, Lee S, Leaf DS, Lily LS, Jordt H, JOhnson T. 2019. A short, course-based research module provides metacognitive benefits in the form of more sophisticated problem solving. Journal of College Science Teaching. 48(4):22–30.

DeHaven B, Sato B, Mello J, Hill T, Syed J, Patel R. 2022. Bootleg biology: A semester-long cure using wild yeast to brew beer. Journal of Microbiology & Biology Education. 23(3).

Deka L, Shereen P, Wand J. 2023. A course-based undergraduate research experience (cure) pathway model in mathematics. PRIMUS. 33:65–83.

Delventhal R, Steinhauer J. 2020. A course-based undergraduate research experience examining neurodegeneration in drosophila melanogaster teaches students to think, communicate, and perform like scientists. PLOS ONE. 15(4):e0230912.

Donegan NT, Zachariah JM, Olimpo JT. 2022. Integrating museum education into an introductory biology cure leads to positive perceptions of scientific research and museum exhibitions among students, faculty, and staff. Journal of Biological Education.

Elkins K, Zeller C. 2020. What is the cure for limited DNA? A forensic science course focused on ngs. The Journal of Forensic Science Education. 2(2).

Ero-Tolliver I, Dumas J. 2019. Work in progress: Students' exposure to cures: Assessing science identity development of underrepresented engineering students at an hbcu. American Society for Engineering Education.

Esparza D, Wagler AE, Olimpo JT. 2020. Characterization of instructor and student behaviors in cure and non-cure learning environments: Impacts on student motivation, science identity development, and perceptions of the laboratory experience. CBE-Life Sciences Education. 19(1):ar10.

Evans CJ, Olson JM, Mondal BC, Kandimalla P, Abbasi A, Abdusamad MM, Acosta O, Ainsworth JA, Akram HM, Albert RB et al. 2021. A functional genomics screen identifying blood cell development genes in drosophila by undergraduates participating in a course-based research experience. G3 (Bethesda). 11(1):jkaa028.

Fornsaglio JL, Sheffler Z, Hull DC. 2021. The impact of semester-long authentic research on student experiences. Journal of Biological Education. 55(1):2–16.

Freeman S, Mukerji J, Sievers M. 2023. A cure on the evolution of antibiotic resistance in escherichia coli improves student conceptual understanding. CBE-Life Sciences Education. 22:ar7.

Furrow RE, Kim HG, Abdelrazek SMR, Dahlhausen K, Yao AI, Eisen JA, Goldman MS, Albeck JG, Facciotti MT. 2020. Combining microbial culturing with mathematical modeling in an introductory course-based undergraduate research experience. Frontiers in Microbiology. 11:581903.

Gasper BJ, Gardner SM. 2013. Engaging students in authentic microbiology research in an introductory biology laboratory course is correlated with gains in student understanding of the nature of authentic research and critical thinking. Journal of Microbiology & Biology Education. 14(1):25–34.

Genet KS. 2021. The cure for introductory, large enrollment, and online courses. SPUR-Scholarship and Practice of Undergraduate Research. 4(3):13–21.

Gerringer ME, Ismail Y, Cannon KA. 2023. Deep-sea biology in undergraduate classrooms: Open access data from remotely operated vehicles provide impactful research experiences. Front Mar Sci. 9:1033274.

Godin EA, Wormington SV, Perez T, Barger MM, Snyder KE, Richman LS, Schwartz-Bloom R, Linnenbrink-Garcia L. 2015. A pharmacology-based enrichment program for undergraduates promotes interest in science. CBE-Life Sciences Education. 14(4):ar40.

Gray C, Price CW, Lee CT, Dewald AH, Cline MA, McAnany CE, Columbus L, Mura C. 2015. Known structure, unknown function: An inquiry-based undergraduate biochemistry laboratory course. Biochemistry and molecular biology education. 43(4):245–262.

Hanauer DI, Graham MJ, Sea P, Betancur L, Bobrownicki A, Cresawn SG, Garlena RA, Jacobs-Sera D, Kaufmann N, Pope WH et al. 2017. An inclusive research education community (irec): Impact of the sea-phages program on research outcomes and student learning. Proceedings of the National Academy of Sciences. 114(51):13531–13536.

Hanauer DI, Nicholes J, Liao FY, Beasley A, Henter H. 2018. Short-term research experience (sre) in the traditional lab: Qualitative and quantitative data on outcomes. CBE-Life Sciences Education. 17(4):ar64.

Hanson PK, Stultz LK. 2022. Linking chemistry and biology through course-based undergraduate research on anticancer ruthenium complexes. Journal of Chemical Education. 99(2):619–628.

Harris D, Schlueter-Kuck K, Austin E. 2021. Course-based undergraduate research in upper-level engineering electives: A case study. Journal of STEM Education. 22(2).

Harvey PA, Wall C, Luckey SW, Langer S, Leinwand LA. 2014. The python project: A unique model for extending research opportunities to undergraduate students. CBE-Life Sciences Education. 13(4):698–710.

Haskew-Layton RE, Minkler JR. 2020. Chick embryonic primary astrocyte cultures provide an effective and scalable model for authentic research in a laboratory class. Journal of Undergraduate Neuroscience Education. 18(2):A86–a92.

Hiatt AC, Hove AA, Ward JR, Ventura L, Neufeld HS, Boyd AE, Clarke HD, Horton JL, Murrell ZE. 2021. Authentic research in the classroom increases appreciation for plants in undergraduate biology students. Integrative and comparative biology. 61(3):969–980.

Hurst-Kennedy J, Saum M, Achat-Mendes C, D'Costa A, Javazon E, Katzman S, Ricks E, Barrera A. 2020. The impact of a semester-long, cell culture and fluorescence microscopy cure on learning and attitudes in an underrepresented stem student population. Journal of Microbiology & Biology Education. 21(1).

Indorf JL, Weremijewicz J, Janos DP, Gaines MS. 2019. Adding authenticity to inquiry in a first-year, research-based, biology laboratory course. CBE-Life Sciences Education. 18(3):ar38.

Irby SM, Pelaez NJ, Anderson TR. 2020. Student perceptions of their gains in course-based undergraduate research abilities identified as the anticipated learning outcomes for a biochemistry cure. Journal of Chemical Education. 97(1):56–65.

Johnson KC, Sabel JL, Cole J, Pruett CL, Plymale R, Reyna NS. 2022. From genetics to biotechnology: Synthetic biology as a flexible course-embedded research experience. Biochemistry and molecular biology education. 50(6):580–591.

Jones CK, Lerner AB. 2019. Implementing a course-based undergraduate research experience to grow the quantity and quality of undergraduate research in an animal science curriculum. Journal of Animal Science. 97(11):4691–4697.

Jordan TC, Burnett SH, Carson S, Caruso SM, Clase K, DeJong RJ, Dennehy JJ, Denver DR, Dunbar D, Elgin SC. 2014. A broadly implementable research course in phage discovery and genomics for first-year undergraduate students. MBio. 5(1):e01051–01013.

Jurgensen SK, Harsh J, Herrick JB. 2021. A cure for salmonella: A laboratory course in pathogen microbiology and genomics. CourseSource.

Kappler U, Rowland SL, Pedwell RK. 2017. A unique large-scale undergraduate research experience in molecular systems biology for non-mathematics majors. Biochemistry and molecular biology education. 45(3):235–248.

Killion PJ, Page IB, Yu V. 2019. Big-data analysis and visualization as research methods for a large-scale undergraduate research program at a research university. SPUR-Scholarship and Practice of Undergraduate Research. 2(4):14–22.

Kinner D, Lord M. 2018. Student-perceived gains in collaborative, course-based undergraduate research experiences in the geosciences. National Science Teachers Association. 48(2):48–58.

Kowalski JR, Hoops GC, Johnson RJ. 2016. "Implementation of a collaborative series of classroom-based undergraduate research experiences spanning chemical biology, biochemistry, and neurobiology". CBE-Life Sciences Education. 15(4).

LaForge J, Martin EC. 2022. Impact of authentic course-based undergraduate research experiences (cures) on student understanding in introductory biology laboratory courses. American Biology Teacher. 84(3):137–142.

Large DN, Van Doorn NA, Timmons SC. 2023. Cancer and chemicals: A research-inspired laboratory exercise based on the ames test for mutagenicity. Biochemistry and molecular biology education. 51(1):103–113.

Laungani R, Tanner C, Brooks TD, Clement B, Clouse M, Doyle E, Dworak S, Elder B, Marley K, Schofield B. 2018. Finding some good in an invasive species: Introduction and assessment of a novel cure to improve experimental design in undergraduate biology classrooms. Journal of Microbiology & Biology Education. 19(2).

Li B, Jia XT, Chi YX, Liu XL, Jia BL. 2020. Project-based learning in a collaborative group can enhance student skill and ability in the biochemical laboratory: A case study. Journal of Biological Education. 54(4):404–418.

Liu JJ, Cook R, Danhof L, Lopatto D, Stoltzfus JR, Benning C. 2021. Connecting research and teaching introductory cell and molecular biology using an arabidopsis mutant screen. Biochemistry and molecular biology education. 49(6):926–934.

Liu Y. 2022. Effects of a cure laboratory module on general chemistry students' perceptions of scientific research, green chemistry, and self-efficacy. Journal of Chemical Education. 99(7):2588–2596.

Lloyd SA, Shanks RA, Lopatto D. 2019. Perceived student benefits of an undergraduate physiological psychology laboratory course. Teaching of Psychology. 46(3):215–222.

Lo SM, Le BD. 2021. Student outcomes from a large-enrollment introductory course-based undergraduate research experience on soil microbiomes. Frontiers in Microbiology. 12:589487.

Lopatto D, Rosenwald AG, Burgess RC, Silver Key C, Van Stry M, Wawersik M, DiAngelo JR, Hark AT, Skerritt M, Allen AK et al. 2022. Student attitudes contribute to the effectiveness of a genomics cure. Journal of Microbiology & Biology Education. 23(2).

Makarevitch I, Frechette C, Wiatros N. 2015. Authentic research experience and "big data" analysis in the classroom: Maize response to abiotic stress. CBE-Life Sciences Education. 14(3).

Malotky MKH, Mayes KM, Price KM, Smith G, Mann SN, Guinyard MW, Veale S, Ksor V, Siu L, Mlo H et al. 2020. Fostering inclusion through an interinstitutional, community-engaged, course-based undergraduate research experience. Journal of Microbiology & Biology Education. 21(1).

Martin A, Rechs A, Landerholm T, McDonald K. 2021. Course-based undergraduate research experiences spanning two semesters of biology impact student self-efficacy but not future goals. Journal of College Science Teaching. 50(4):33–47.

May NW, McNamara SM, Wang S, Kolesar KR, Vernon J, Wolfe JP, Goldberg D, Pratt KA. 2018. Polar plunge: Semester-long snow chemistry research in the general chemistry laboratory. Journal of Chemical Education. 95(4):543–552.

Mayer B, Blume A, Black C, Stevens S. 2019. Improving student learning outcomes through community-based research: The poverty workshop. Teach Sociol. 47(2):135–147.

McDonough J, Goudsouzian LK, Papaj A, Maceli AR, Klepac-Ceraj V, Peterson CN. 2017. Stressing escherichia coli to educate students about research: A cure to investigate multiple levels of gene regulation. Biochemistry and molecular biology education. 45(5):449–458.

McLaughlin JS. 2021a. Teaching environmental sustainability while transforming study abroad. Sustainability-Basel. 13(1).

McLaughlin JS, Patel M, Slee JB. 2020. A cure using cell culture-based research enhances career-ready skills in undergraduates. SPUR-Scholarship and Practice of Undergraduate Research. 4(2):49–61.

McLaughlin KJ. 2021b. Developing a macromolecular crystallography driven cure. Structural Dynamics. 8:020406.

Miller CW, Hamel J, Holmes KD, Helmey-Hartman WL, Lopatto D. 2013. Extending your research team: Learning benefits when a laboratory partners with a classroom. Bioscience. 63(9):754–762.

Mordacq JC, Drane DL, Swarat SL, Lo SM. 2017. Development of course-based undergraduate research experiences using a design-based approach. Journal of College Science Teaching. 46(4):64–75.

Nadelson L, Walters L, Waterman J. 2010. Course-integrated undergraduate research experiences structured at different levels of inquiry. Journal of STEM Education. 11(1).

Ochoa SD, Dores MR, Allen JM, Tran T, Osman M, Castellanos NPV, Trejo J, Zayas RM. 2019. A modular laboratory course using planarians to study genes involved in tissue regeneration. Biochemistry and molecular biology education. 47(5):547–559.

Olimpo JT, Fisher GR, DeChenne-Peters SE. 2016. Development and evaluation of the tigriopus course-based undergraduate research experience: Impacts on students' content knowledge, attitudes, and motivation in a majors introductory biology course. CBE-Life Sciences Education. 15(4).

Olson JM, Evans CJ, Ngo KT, Kim HJ, Nguyen JD, Gurley KGH, Ta T, Patel V, Han L, Truong NK et al. 2019. Expression-based cell lineage analysis in drosophila through a course-based research experience for early undergraduates. G3 (Bethesda). 9(11):3791–3800.

Ott LE, Godsay S, Stolle-McAllister K, Kowalewski C, Maton KI, LaCourse WR. 2020. Introduction to research: A scalable, online badge implemented in conjunction with a classroom-based undergraduate research experience (cure) that. UI J. 11(1).

Overath R, Zhang D, Hatheril J. 2016. Implementing course-based research increases student aspirations for stem degrees. Council on Undergraduate Research Quarterly. 37(2).

Pagano JK, Jaworski L, Lopatto D, Waterman R. 2018. An inorganic chemistry laboratory course as research. Journal of Chemical Education. 95(9):1520–1525.

Papendieck A, Ellins K, Clarke J. 2020. Developing a disciplinarily diverse course-based research experience: Outcomes and design considerations. ICLS Proceedings. 3:1777–1778.

Pavlova IV, Remington DL, Horton M, Tomlin E, Hens MD, Chen D, Willse J, Schug MD. 2021. An introductory biology research-rich laboratory course shows improvements in students' research skills, confidence, and attitudes. PLOS ONE. 16(12).

Pedwell RK, Fraser JA, Wang JTH, Clegg JK, Chartres JD, Rowland SL. 2018. The beer and biofuels laboratory: A report on implementing and supporting a large, interdisciplinary, yeast-focused course-based undergraduate research experience. Biochemistry and molecular biology education. 46(3):213–222.

Penner MR, Sathy V, Hogan KA. 2021. Inclusion in neuroscience through high impact courses. Neuroscience Letters. 750.

Peteroy-Kelly MA, Marcello MR, Crispo E, Buraei Z, Strahs D, Isaacson M, Jaworski L, Lopatto D, Zuzga D. 2017. Participation in a year-long cure embedded into major core genetics and cellular and molecular biology laboratory courses results in gains in foundational biological concepts and experimental design skills by novice undergraduate researchers. Journal of Microbiology & Biology Education. 18(1).

Peterson CN, Callahan KP. 2022. External collaboration results in student learning gains and positive stem attitudes in cures. CBE-Life Sciences Education. 21(4):ar74.

Pieczynski JN, Deets A, McDuffee A, Lynn Kee H. 2019. An undergraduate laboratory experience using crispr-cas9 technology to deactivate green fluorescent protein expression in escherichia coli. Biochemistry and molecular biology education. 47(2):145–155.

Ramirez-Lugo JS, Toledo-Hernandez C, Velez-Gonzalez I, Ruiz-Diaz CP. 2021. Creare: A course-based undergraduate research experience to study the responses of the endangered coral acropora cervicornis to a changing environment. Journal of Microbiology & Biology Education. 22(1).

Reed KE, Richardson JM. 2012. Using microbial genome annotation as a foundation for collaborative student research. Biochemistry and molecular biology education. 41(1).

Roberts LA, Shell SS. 2022. A research program-linked, course-based undergraduate research experience that allows undergraduates to participate in current research on mycobacterial gene regulation. Frontiers in Microbiology. 13:1025250.

Rubush DM, Stone KL. 2020. A learning community involving collaborative course-based research experiences for foundational chemistry laboratories. Education Sciences. 10(4).

Russell JE, D'Costa A, Runck C, Barnes D, Barrera A, Hurst-Kennedy J, Sudduth E, L. QE, Schlueter M, Iskhakova L et al. 2015. Bridging the undergraduate curriculum using an integrated course-embedded undergraduate research experience (icure). CBE-Life Sciences Education. 14(1):ar4.

Ruth A, Brewis A, SturtzSreetharan C. 2023. Effectiveness of social science research opportunities: A study of course-based undergraduate research experiences (cures). Teaching in Higher Education. 28(7):1484–1502.

Saha A, Williams L. 2021. Engaging undergraduate students in authentic research in the inorganic chemistry laboratory course. Engaged Student Learning: Essays on Best Practices in the University System of Georgia 3.

Sandquist EJ, Cervato C, Ogilvie C. 2019. Positive affective and behavioral gains of first-year students in course-based research across disciplines. SPUR-Scholarship and Practice of Undergraduate Research. 2(4):45–57.

Sarmah S, Chism GW, Vaughan MA, Muralidharan P, Marrs JA, Marrs KA. 2016. Using zebrafish to implement a course-based undergraduate research experience to study teratogenesis in two biology laboratory courses. Zebrafish. 13(4):293–304.

Satusky MJ, Wilkins H, Hutson B, Nasiri M, King DE, Erie DA, Freeman TC. 2022. Cureing biochemistry lab monotony. Journal of Chemical Education. 99(12):3888–3898.

Sewall JM, Oliver A, Denaro K, Chase AB, Weihe C, Lay M, Martiny JBH, Whiteson K. 2020. Fiber force: A fiber diet intervention in an advanced course-based undergraduate research experience (cure) course. Journal of Microbiology & Biology Education. 21(1).

Shaffer CD, Alvarez CJ, Bednarski AE, Dunbar D, Goodman AL, Reinke C, Rosenwald AG, Wolyniak MJ, Bailey C, Barnard D et al. 2014. A course-based research experience: How benefits change with increased investment in instructional time. CBE-Life Sciences Education. 13(1):111–130.

Shaner SE, Hooker PD, Nickel AM, Leichtfuss AR, Adams CS, de la Cerda D, She YQ, Gerken JB, Pokhrel R, Ambrose NJ et al. 2016. Discovering inexpensive, effective catalysts for solar energy conversion: An authentic research laboratory experience. Journal of Chemical Education. 93(4):650–657.

Sharma AK, Hernandez M, Phuong V. 2019. Engaging students with computing and climate change through a course in scientific computing. Journal of STEM Education. 20(2):49–57.

Shuster MI, Curtiss J, Wright TF, Champion C, Sharifi M, Bosland J. 2019. Implementing and evaluating a course-based undergraduate research experience (cure) at a hispanic-serving institution. Interdisciplinary Journal of Problem-Based Learning. 13(2).

Smith KPW, Waddell EA, Dean AN, Anandan S, Gurney S, Kabnick K, Little J, McDonald M, Mohan J, Marenda DR et al. 2023. Course-based undergraduate research experiences are a viable approach to increase access to research experiences in biology. Journal of Biological Education. 57(3):618–632.

Smith MA, Olimpo JT, Santillan KA, McLaughlin JS. 2022. Addressing foodborne illness in cote d'ivoire: Connecting the classroom to the community through a nonmajors course-based undergraduate research experience. Journal of Microbiology & Biology Education. 23(1).

Smyth DS. 2018. An authentic course-based research experience in antibiotic resistance and microbial genomics. Science education and civic engagement. 9(2).

Sorensen AE, Corral L, Dauer JM, Fontaine JJ. 2018. Integrating authentic scientific research in a conservation course-based undergraduate research experience. Natural Sciences Education. 47(1):1–10.

Spence NJ, Anderson R, Corrow S, Dumais SA, Dierker L. 2022. Passion-driven statistics: A course-based undergraduate research experience (cure). Mathematics Enthusiast. 19(3):759–770.

Stanfield E, Slown CD, Sedlacek Q, Worcester SE. 2022. A course-based undergraduate research experience (cure) in biology: Developing systems thinking through field experiences in restoration ecology. CBE-Life Sciences Education. 21(2):ar20.

Staub NL, Poxleitner M, Braley A, Smith-Flores H, Pribbenow CM, Jaworski L, Lopatto D, Anders KR. 2016. Scaling up: Adapting a phage-hunting course to increase participation of first-year students in research. CBE-Life Sciences Education. 15(2).

Stoeckman AK, Cai Y, Chapman KD. 2019. Icure (iterative course-based undergraduate research experience): A case-study. Biochemistry and molecular biology education. 47(5):565–572.

Stovall GM, Huynh V, Engelman S, Ellington AD. 2019. Aptamers in education: Undergraduates make aptamers and acquire 21st century skills along the way. Sensors. 19(15).

Strain AK, Williams MA, Phillips J. 2021. Modeling the research process: Authentic human physiology research in a large non-majors course. CourseSource.

Thu YM, French LB, Hersh BM, Nelson MK, Whitenack LB. 2021. Students' perceptions of fsbio 201, a cure-based course that scaffolds research and scientific communication, align with learning outcomes. Integrative and comparative biology. 61(3):944–956.

Tomasik JH, Cottone KE, Heethuis MT, Mueller A. 2013. Development and preliminary impacts of the implementation of an authentic research-based experiment in general chemistry. Journal of Chemical Education. 90(9):1155–1161.

Tomasik JH, LeCaptain D, Murphy S, Martin M, Knight RM, Harke MA, Burke R, Beck K, Acevedo-Polakovich ID. 2014. Island explorations: Discovering effects of environmental research-based lab activities on analytical chemistry students. Journal of Chemical Education. 91(11):1887–1894.

Vater A, Mayoral J, Nunez-Castilla J, Labonte JW, Briggs LA, Gray JJ, Makarevitch I, Rumjahn SM, Siegel JB. 2021. Development of a broadly accessible, computationally guided biochemistry course-based undergraduate research experience. Journal of Chemical Education. 98(2):400–409.

Villa-Cuesta E, Hobbie L. 2016. Genetics research project laboratory: A discovery-based undergraduate research course. Genetics Society of America Peer-Reviewed Education Portal. 3.

Waddell EA, Ruiz-Whalen D, O'Reilly AM, Fried NT. 2021. Flying in the face of adversity: A drosophila-based virtual cure (course-based undergraduate research experience) provides a semester-long authentic research opportunity to the flipped classroom. Journal of Microbiology & Biology Education. 22(3).

Ward JR, Clarke HD, Horton JL. 2014. Effects of a research-infused botanical curriculum on undergraduates' content knowledge, stem competencies, and attitudes toward plant sciences. CBE-Life Sciences Education. 13(3):387–396.

Werth A, West CG, Lewandowski HJ. 2022. Impacts on student learning, confidence, and affect and in a remote, large-enrollment, course-based undergraduate research experience in physics. Physical Review Physics Education Research. 18(1).

Wickham RJ, Genné-Bacon EA, Jacob MH. 2021. The spine lab: A short-duration, fully-remote course-based undergraduate research experience. Journal of Undergraduate Neuroscience Education. 20(1):A28–a39.

Wilczek LA, Clarke AJ, Martinez MDG, Morin JB. 2022. Catalyzing the development of self-efficacy and science identity: A green organic chemistry cure. Journal of Chemical Education. 99(12):3878–3887.

Williams LC, Reddish MJ. 2018. Integrating primary research into the teaching lab: Benefits and impacts of a one-semester cure for physical chemistry. Journal of Chemical Education. 95(6):928–938.

Woodley SK, Freeman PE, Ricketts TD. 2019. Combining novel research and community-engaged learning in an undergraduate physiology laboratory course. Advances in physiology education. 43(2):110–120.

Wooten MM, Coble K, Puckett AW, Rector T. 2018. Investigating introductory astronomy students' perceived impacts from participation in course-based undergraduate research experiences. Physical Review Physics Education Research. 14(1).

Yau YCM, Cheung TSE, Chen XF, Wong CM. 2022. Use of course-based undergraduate research experiences model to enhance research interest of hong kong health professional undergraduate students. 8th International Conference On Higher Education Advances (Head '22).773–779.

Zelaya AJ, Gerardo NM, Blumer LS, Beck CW. 2020. The bean beetle microbiome project: A course-based undergraduate research experience in microbiology. Frontiers in Microbiology. 11:577621.

Zhou YY, Jung E, Arróyave R, Radovic M, Shamberger P. 2015. Incorporating research experiences into an introductory materials science course. Int J Eng Educ. 31(6):1491–1503.
